# Aging impairs skeletal muscle regeneration by promoting fibro/fatty degeneration and inhibiting inflammation resolution via fibro-adipogenic progenitors

**DOI:** 10.1101/2023.11.27.568776

**Authors:** Francisco Garcia-Carrizo, Sabrina Gohlke, Georgia Lenihan-Geels, Anne-Marie Jank, Marina Leer, George A. Soultoukis, Masoome Oveisi, Catrin Herpich, Claudia A. Garrido, Georgios Kotsaris, Sophie Pöhle-Kronawitter, Arnold Tsamo-Tetou, Antonia Graja, Mario Ost, Laura Villacorta, Raphael S. Knecht, Susanne Klaus, Annette Schürmann, Sigmar Stricker, Katharina Schmidt-Bleek, Amaia Cipitria, Georg N. Duda, Vladimir Benes, Ursula Müller-Werdan, Kristina Norman, Tim J. Schulz

## Abstract

Skeletal muscle regeneration depends on the function of fibro/adipogenic progenitors (FAPs). Here we show that aging impairs myogenic stem cells by disrupting the integration of extracellular matrix and immunomodulatory functions within the stem cell niche, thereby promoting fibro/fatty degeneration. We identify the FAP-secreted protein Periostin as a niche factor that is decreased in aged muscle and in circulation of aged humans with low-exercise lifestyle. Periostin controls FAP-expansion after injury and its depletion fate-regulates FAPs towards adipogenesis. This leads to delayed pro- to anti-inflammatory macrophage transition during regeneration. Transplantation of young FAPs with high Periostin secretion, but not Periostin-deficient FAPs, into aged muscle restores inflammation resolution and successful regeneration. Mechanistically, Periostin activates Focal adhesion kinase- and AKT-signaling in macrophages via integrins to promote an anti-inflammatory profile, which synchronizes matrix-derived mechanosensory signaling and immunomodulation. These results uncover a novel role of FAP-based regulation that orchestrates successful muscle regeneration and prevents fibro/fatty degeneration.

## Introduction

Aging is characterized by a gradual loss of muscle mass and strength that leads to sarcopenia as part of the frailty syndrome, a pathology that constitutes a serious health problem associated with adverse outcomes in elderly individuals^1,2^. This age-related decline in muscle function is not only associated with a reduction in muscle stem cell (MuSC) numbers and a reduced regenerative potential but also the accumulation of fibrotic tissue and ectopic adipocytes within the skeletal muscle, which is referred to as fibro/fatty infiltration or fatty degeneration^3–7^. MuSCs also interact with their local microenvironment and a key component of the muscle stem cell niche is the extracellular matrix (ECM) which experiences extensive remodeling during regeneration and throughout life^8–12^. Fibro/adipogenic progenitor cells (FAPs) are considered the main ECM-source, they regulate its remodeling and are indispensable for long-term muscle maintenance and MuSCs’ functionality during regeneration^13–15^. However, under pathological conditions, such as muscular dystrophies, obesity, and type 2 diabetes mellitus (T2DM), subsets of FAPs are thought to generate fibrotic scar tissue and differentiate into adipocytes that progressively replace functional muscle tissue, resulting in fibro/fatty degeneration and the age-related metabolic dysfunction of muscle tissue^6,16–20^.

Besides their role in ECM remodeling, FAPs secrete trophic factors that control MuSC activation through different mechanisms^14,21–23^ and aged FAPs display an aberrant secretory profile of WISP1, BMP3b or SMOC2, among others, that reduce MuSCs’ regeneration capacity^10,11,24^. The contribution of aging to FAPs’ dysregulation and fibro/fatty infiltration in skeletal muscle remains less well understood. In addition to the FAP-MuSCs niche crosstalk, FAPs interact with different immune cells^25–28^. For instance, pro-inflammatory macrophages regulate the number of FAPs via TNF-mediated apoptosis^28^. A failure to resolve the initial pro-inflammatory macrophage response, however, reduces myogenic regeneration and promotes chronic inflammation in aged muscles^29–32^.

Periostin (*Postn*) is a matricellular protein that interacts with ECM components to drive fibrillogenesis and matrix scaffold formation. It also displays signaling activity by binding integrin receptors, mainly αvβ3 and αvβ5 heterodimers, to promote cell adhesion, migration and proliferation in different tissues and in the context of regeneration^33–35^. Expression of *Postn* is transiently induced in FAPs during muscle regeneration and initial data suggest a role in regeneration by coupling myogenesis and angiogenesis^36–38^. Omics-based analyses also list *Postn* among the potentially dysregulated genes in aged muscles^11,22^. In addition to its tissue/stem cell-specific function, Periostin is discussed as a circulating biomarker of type-2 inflammation in asthma, breast cancer, cardiovascular disease and liver steatosis^39–42^, as well as a potential biomarker of aging in serum and across several murine tissues^22,43^.

We here investigate the role of age-related changes of the muscle stem cell nice, which result in increased fibro/fatty degeneration of skeletal muscle at the expense of healthy tissue regeneration. We identify a mechanism that synchronizes several key ECM-functions, including mechano-signaling and immunomodulation, between the different cell populations of the muscle microenvironment to orchestrate successful muscle regeneration. This niche factor-mediated integration is impaired during aging and we identify the matricellular protein Periostin as an important contributor to these processes. Loss of Periostin phenocopies muscle aging, resulting in delayed regeneration, ectopic adipocyte accumulation and fibrosis, which contribute to delayed inflammation resolution during injury repair. Reconstitution of Periostin through FAP-transplantation recovers young-like regeneration in aged muscle. Mechanistically, Periostin targets mechanosensory pathways in niche-resident macrophages, thereby promoting resolution of the inflammatory microenvironment during regeneration. Periostin may serve as sarcopenia biomarker as circulating levels positively correlate with muscle mass and function at different ages, but are reduced with age in humans.

## Results

### Loss of Periostin in aged muscle disrupts extracellular matrix function and the stem cell niche

To examine the contribution of aging to myogenic niche disruption, we compared young, 2-months-old male and female mice to sex-matched animals aged 15-18 months. This age was chosen to reduce the likelihood of other, potentially confounding, pathologies that may occur in even older mice. At 15 months, muscles display a gene expression profile and morphology consistent with fibro/fatty degeneration, suggesting such changes are relatively early hallmarks of muscle aging (Figures 1A, S1A-S1C). Fluorescence-activated cell sorting (FACS) showed increased numbers of muscle-resident FAPs with age while MuSCs decreased in male and female mice (Figures 1B, S1D-S1H). Primary FAPs isolated from muscle of 15-months-old mice displayed spontaneous lipid droplet accumulation, while aged MuSCs showed reduced myogenic potential compared to respective young-muscle cell cultures (Figures 1C and 1D). Given the marked increase in adipogenic potential in aged FAPs, we decided to further examine this cell population. Microarray analysis of freshly sorted FAPs showed a marked reduction in many genes and pathways linked to the ECM (Table S1, Figure S1I). The top down-regulated gene, *Postn*, which encodes for Periostin, is a secreted matricellular protein^35^ and could thus act as paracrine niche factor (Figure 1E). Aging muscles displayed reduced Periostin expression on protein and mRNA levels in male and female mice, and cultured young FAPs readily released Periostin into the supernatant, while aged FAPs only secreted minor amounts (Figures 1F-1H, S1J and S1K).

**Figure 1:**
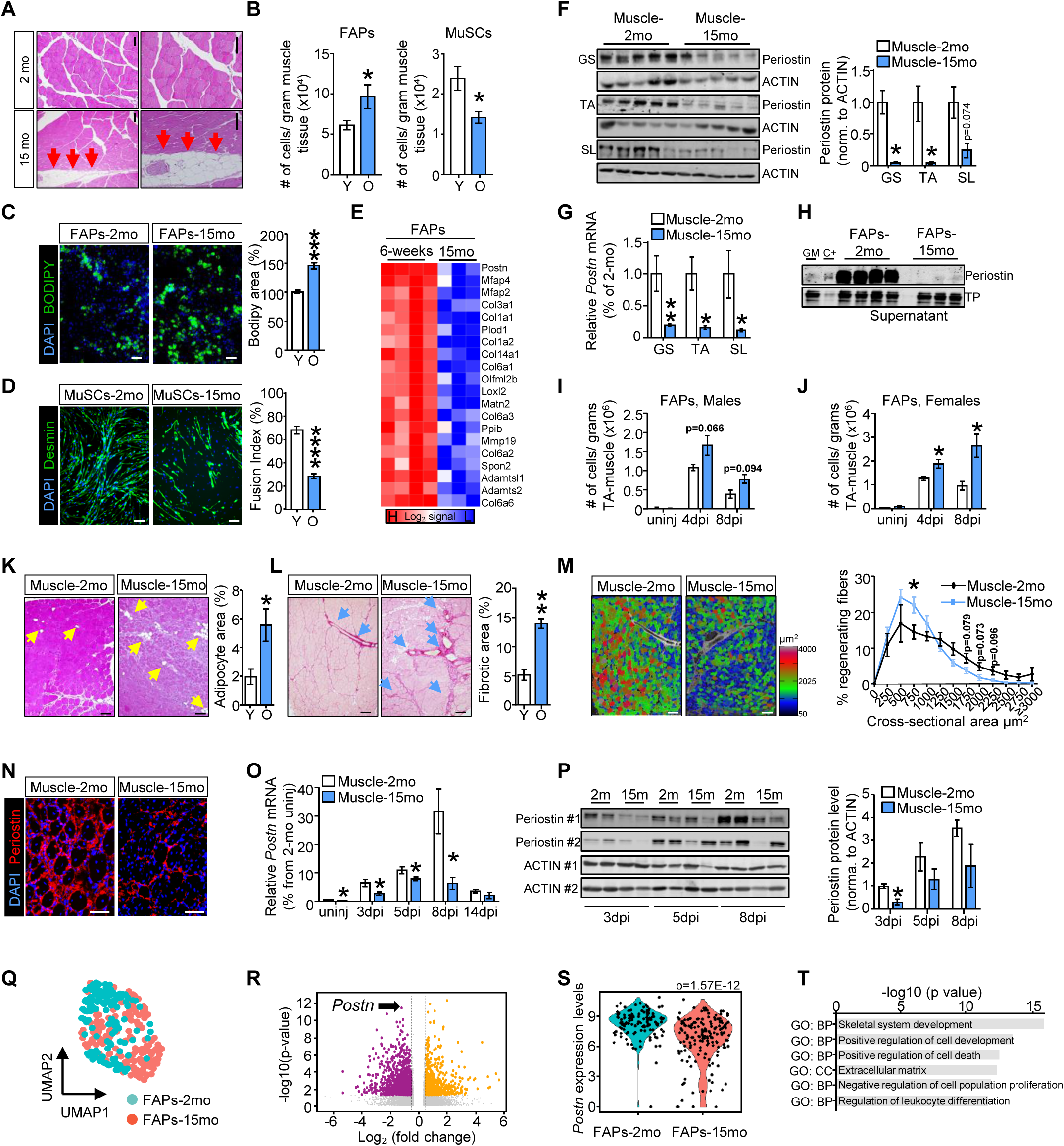
Aging impairs myogenic regeneration by disrupting FAPs-derived niche signals. (**A**) Representative hematoxylin/eosin staining images of gastrocnemius muscle cross-sections showing inter-myocellular adipocyte infiltration between myofibers from young (2 months) and aged (age: 15-18 months; pertains to all subsequent panels) male mice (red arrows indicate adipocytes; scale bar: 100 µm). (**B**) Quantifications of flow cytometric analysis of CD45^-^CD31^-^SCA1^+^ FAPs and CD45^-^CD31^-^SCA1^-^ITGA7^+^ MuSCs per grams of murine muscle tissue (n=18/group). White bars indicate young mice/cells (age: 2-2.5 months), blue bars indicate aged mice/cells (age: 15-18 months; pertains to all subsequent panels). (**C**) Representative images and quantification of spontaneous adipogenic differentiation of FAPs (green: BODIPY, lipid droplet; blue: DAPI, nuclei; n=6; 2 independent experiments; scale bar: 50 µm). (**D**) Representative images and fusion index quantifications in differentiated MuSC (green: Desmin, myofibers; blue: DAPI-nuclei; n=6; 2 independent experiments; scale bar, 100 µm). (**E**) Heat map of top-20 down-regulated ECM genes in microarray analysis comparing freshly sorted FAPs from uninjured skeletal muscle. (**F**) Periostin protein detection and quantification by western blot (Actin: loading control; n=5/group). (**G**) *Postn* mRNA expression in uninjured gastrocnemius (GS), tibialis anterior (TA) and soleus (SL) muscles (n=5/group). (**H**) Periostin protein detection by western blot of cell culture supernatants of FAPs collected during adipogenic differentiation 48h after last medium change (C+: mouse rPOSTN; TP: total protein loading control). (**I**, **J**) FACS-analysis of CD45^-^ CD31^-^SCA1^+^ FAPs per grams of TA muscle comparing uninjured to injured TA at 4- and 8-dpi (n=3/group for uninjured and 8-dpi; n=6/group for 4-dpi). (**K**) Representative hematoxylin/eosin staining and quantification of adipocyte infiltration in injury sites of TA at 14-dpi (yellow arrows indicate adipocyte areas; n=3/group; scale bar: 50 µm). (**L**) Representative Sirius red staining and quantification of fibrotic tissue deposition in TA injury sites at 14-dpi (blue arrows indicate fibrosis areas; n=3/group; scale bar: 50 µm). (**M**) Representative images and CSA quantification of fiber size ranges in injury sites at 14-dpi (n=4/group; scale bar: 100 µm). (**N**) Representative images of Periostin protein immunofluorescence in injury sites in TA at 8-dpi (red: Periostin; blue: DAPI, nuclei; scale bar: 50 µm). (**O**) *Postn* mRNA levels in uninjured TA-muscles compared to TA at 3-, 5-, 8- and 14-dpi (n=4/group). (**P**) Periostin protein western blots and quantification (Actin: loading control) in TA at 3-, 5-, and 8-dpi (n=4/group). (**Q**) UMAP dimensionality reduction of scRNA-seq analysis of 287 CD45^-^CD31^-^SCA1^+^ FAPs FACS-isolated at 4-dpi from young (FAPS-2mo, 128 cells; blue) and aged (FAPs-15mo, 159 cells; red) mice generated using microfluidics-based technology, and (**R**) volcano plot of DEGs (p<0.05 and |Log2|fold change ≥0.5 as cutoffs; black arrow indicates *Postn*). (**S**) Violin plot depiction of *Postn* gene expression per cell in scRNA-seq dataset. (**T**) Enriched Gene Ontology terms of biological process (BP) and cellular component (CC) of downregulated DEGs from scRNA-seq dataset. Data were collected from male mice except in (J, female). All data are presented as mean ± standard error of the mean (SEM), statistical significances are *p<0.05, **p<0.01, ***p<0.001.

We next tested whether muscle injury would elicit a similar phenotype and applied an established model of tibialis anterior (TA) muscle injury by intramuscular injection of glycerol, a protocol that induces ectopic adipogenesis and fibrosis at the injury site^44^. Injury-induced expansion of FAPs was more pronounced in aged TA compared to young post-injury muscle in male and female mice. Unexpectedly, MuSC expansion was also induced in aged muscle of female mice and to some extent in male mice Figures 1I, 1J and S1L-S1N). Clearance of FAPs is critical to prevent fibro/fatty infiltration^28^, and accordingly we detected increased adipogenesis and fibrosis areas in injured TA of aged compared to young male mice which was accompanied by an age-related reduction in myofiber cross-sectional areas (CSA) at the injury site and reduced expression of myogenesis markers (Figures 1K-1M and S1O-S1Q). To further explore the involvement of Periostin in muscle healing, we evaluated its expression profile on multiple days post-injury (dpi). In line with other publications^36–38,45^, we observed an increase of Periostin in the interstitial space between regenerating fibers and corresponding dynamic mRNA and protein expression patterns that peaked between 5- and 8- dpi in muscle of young mice, all of which was markedly attenuated in aged muscle (Figures 1N-1P). We verified disrupted FAP-functions in regenerating aged muscles by applying single cell-RNA sequencing (scRNA-seq) of FAPs isolated from 4-dpi muscle. Uniform manifold approximation and projection (UMAP) visualization and differentially expressed gene (DEG) analysis revealed partial transcriptional separation of FAPs of the two age groups (Figure 1Q; Table S2). This analysis confirmed *Postn* as one of the most significantly down-regulated genes and the enrichment of ECM-related pathways when assessing all significantly downregulated genes (Figures 1R-1T). Up-regulated pathways and a subsequent analysis of predicted secretome genes showed a potential impact of FAP-aging on the modulation of inflammatory responses (Figures S1R and S1S). These findings taken together indicate that aging negatively influences niche functions of FAPs which contribute to fibro/fatty degeneration of aged muscles.

### Genetic inactivation of *Postn* recapitulates an aging-like phenotype and promotes fibro/fatty degeneration

To determine whether loss of Periostin could indeed contribute to a decline of muscle regeneration as observed in aged mice, we obtained a mouse model with genetic inactivation of the *Postn* gene by insertion of a LacZ cassette, leading to loss of the *Postn* gene product and expression of a β-galactosidase reporter under control of the native *Postn* promoter (Figures S2A-S2D). Periostin protein was completely absent in muscles of *Postn^-/-^* mice whereas the reporter was expressed in the interstitial space between myofibers, consistent with the localization of FAPs (Figure S2B). No differences in myofiber CSAs or frequencies of FAPs and MuSCs were observed in 2-3 months-old muscles of male or female *Postn^-/-^* mice without injury. However, we noticed that dissected muscles displayed a significant reduction in mechanical stiffness (Figures S2E to S2G). Similar to aged muscle, isolated *Postn^-/-^*-FAPs displayed significantly enhanced spontaneous adipogenesis in culture whereas cultured MuSCs had significantly lower myogenic potential (Figures 2A and 2B). Applying the glycerol-injection regeneration model to *Postn^-/-^* mice and littermate wildtype controls using both male and female offspring showed a phenotype closely resembling the phenotype seen in aged muscle: We consistently observed increased expansion of FAPs and MuSCs in *Postn^-/-^* mice (Figure 2C, 2D, S2H and S2I). Importantly, both sexes showed impaired regeneration in *Postn^-/-^* mice with significant accumulation of adipocytes and fibrotic structures and reduced CSA of newly regenerated myofibers at 14-dpi, which was also reflected by increased expression of adipocyte and fibrosis genes and reduced expression of myogenesis markers (Figures 2E-2G and S2J-S2Q). In summary, genetic ablation of *Postn* recapitulates an aging-like disruption of regeneration in male and female mice at the level of FAPs and MuSCs, resulting in fibro/fatty degeneration.

**Figure 2:**
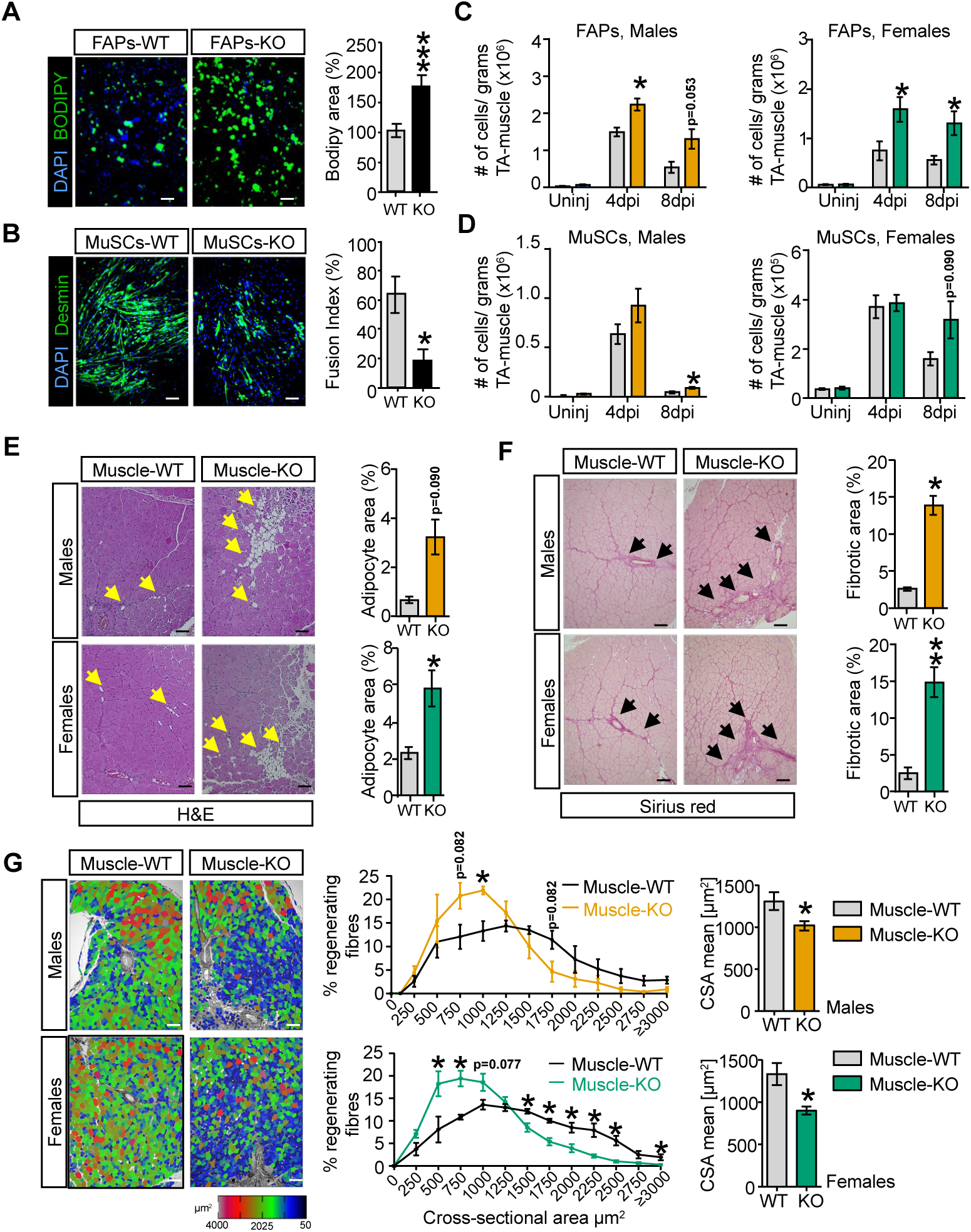
Ablation of *Postn* gene recapitulates an aging-like phenotype in muscle. (**A**) Representative images and quantification of spontaneous adipogenic differentiation of FAPs from *Postn*-KO (black bar) compared to wildtype (WT) littermates (gray bar; green: BODIPY, lipid droplet; blue: DAPI, nuclei; n=8/group; 2 independent experiments; scale bar: 50 µm). (**B**) Representative images and fusion index quantification in differentiated MuSC from *Postn*^-/-^ (KO; black bar) compared to WT littermates (gray bar; green: Desmin, myofibers; blue: DAPI, nuclei; n=3/group; scale bar: 100 µm). (**C**, **D**) Flow cytometric analysis of CD45^-^CD31^-^SCA1^+^ FAPs (panel C) and CD45^-^CD31^-^SCA1^-^ ITGA7^+^ MuSCs (panel D) per grams of TA-muscle comparing uninjured to injured TA at 4- and 8-dpi (males: uninj n=4/3, 4-dpi n=4/3, 8-dpi n=4/4; females: uninj n=10/10, 4-dpi n=7/7, 8-dpi n=4/4). Data show comparisons of male WT (gray bars, black line) to KO (orange bars and line) and female WT (gray bars, black line) to KO (green bars and line) mice (also applies to subsequent panels). (**E**) Representative hematoxylin/eosin staining and quantification of adipocyte infiltration in injury sites of TA at 14-dpi (yellow arrows indicate adipocyte areas; males: n=3/group, females: n=6/7/group; scale bar: 50 µm). (**F**) Representative Sirius red staining and quantification of fibrotic tissue deposition in TA injury sites at 14-dpi (black arrows indicate fibrosis areas; males: n=3/group, females: n=3/4/group; scale bar: 50 µm). (**G**) Representative images, CSA quantification of fiber size ranges and average in injury site at 14- dpi (males: n=3/group; females: n=4/group; scale bar, 100 µm). All data are presented as mean ± standard error of the mean (SEM), statistical significances are *p<0.05, **p<0.01, ***p<0.001.

### Periostin rescues stem cell function and muscle regeneration in aged mice

To better understand the function of Periostin as ECM-related niche factor, we tested the impact of a mouse recombinant Periostin (mrPOSTN) on FACS-purified FAPs and MuSCs of aged and *Postn*-deficient mice. Both cell types, aged and *Postn^-/-^*-FAPs, displayed significantly increased proliferation when compared to the respective control FAPs, whereas MuSCs displayed reduced proliferation rates in comparison to respective controls (Figures 3A-3D, S3A and S3B). After 48h of culture in the presence of mrPOSTN the respective FAP- and MuSC-proliferation phenotypes were fully reversed in *Postn^-/-^* and aged cells. Considering that FAPs are likely a main source of muscle Periostin, these data suggest that FAPs auto-inhibit their own expansion while stimulating cell cycle in MuSCs through paracrine Periostin (Figures 1H, 3A-3D). To determine whether FAP-derived Periostin influences the function of MuSCs, we used an in vitro co-culture system. Consistent with our initial observation, lack of Periostin in MuSCs significantly reduced fusion capacity and resulted in shorter myofibers during differentiation (Figures 3E-3G). Co-cultures of wildtype, Periostin-secreting FAPs, but not *Postn^-/-^*-FAPs, with *Postn*-deficient MuSCs resulted in a full recovery of the impaired myogenesis phenotype (Figures 3E-3G). As cultured MuSCs themselves secrete very little Periostin in comparison to FAPs, this interaction appears to be a key element in maintaining the in vitro myogenic potential of the myogenic stem cell pool (Figure S3C).

**Figure 3:**
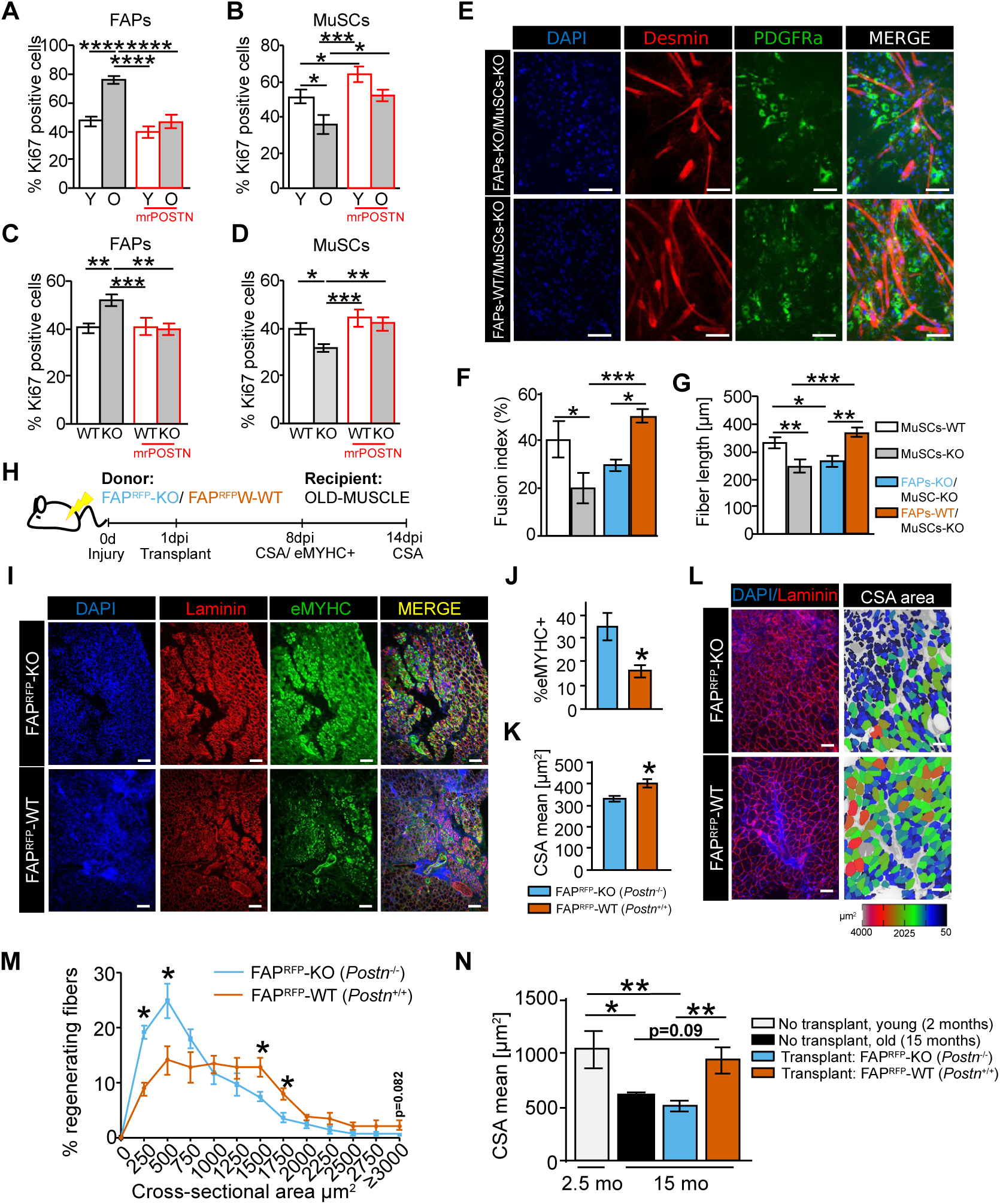
FAPs-secreted Periostin rescues age-related defect of myogenic regeneration. (**A**-**D**) Proliferation analysis by KI67 immunostaining of FAPs and MuSCs FACS-isolated from young (Y, 2 months) and aged (O, 15 months) mice and young *Postn*^-/-^ (KO) and littermate wildtype (WT) control mice and treated with mrPOSTN (0.5 µg/ml) for 48h (n=6-12; two independent experiments). (**E**) Representative images of co-cultures from FAPs-WT and FAPs-KO with MuSCs-KO co-cultured for 5 days in myogenic differentiation medium (green: PDGFRα, FAPs; red: Desmin, myofibers; blue: DAPI, nuclei; scale bar: 50µm). (**F**, **G**) Quantifications of fusion index (% of myofibers with >3 DAPI^+^ nuclei) and myofiber length in co-cultures in panel E, compared to pure MuSC cultures from wildtype (MuSCs-WT) or *Postn*^-/-^ mice (MuSCs-KO) (n=6 wells; two-independent experiments). (**H**) Experimental layout and timeline of FAPs transplants of young FAP^RFP^-WT and FAP^RFP^-KO into old recipient mice (TA-muscle, 15 months). (**I**) Representative immunofluorescence images of transplanted TA-muscle sections at 8-dpi (green: eMHC; red: Laminin; blue: DAPI, nuclei; scale bar: 100 µm). (**J**, **K**) Quantifications of eMHC (top) and CSA average (bottom) of FAP^RFP^-WT and FAP^RFP^-KO transplants at 8-dpi (n=5/group). (**L**, **M**) Representative images and CSA quantification of fiber size ranges in injury site at 14-dpi comparing 15-months TA-muscle transplanted with FAP^RFP^-WT or FAP^RFP^-KO transplants (red: Laminin; blue: DAPI, nuclei; n=5-6/group; scale bar: 100 µm). (**N**) Quantification of CSA averages in regenerating fibers comparing non-transplanted (as shown in Figure 1M) young (2 months; n=4) and old (15 months; n=4) to transplanted 15-months old TA-muscles with either FAP^RFP^-WT (n=5) or FAP^RFP^-KO (n=6). All data are presented as mean ± standard error of the mean (SEM), statistical significances are *p<0.05, **p<0.01, ***p<0.001.

To determine whether Periostin reconstitution has the capacity to enhance the regeneration process in aged skeletal muscle, we sorted FAPs from a mouse model carrying a red fluorescent protein (RFP) expression allele in addition to the *Postn*-knockout allele (Figures S3D and S3E) ^46^. RFP^+^ FAPs either with (FAP^RFP^-WT) or without (FAP^RFP^-KO) *Postn* expression were transplanted into TA muscle of 15-months-old recipient mice one day after glycerol-injury. Transplanted FAPs remained detectable based on RFP expression until at least 8-dpi (Figures 3H, S3D-S3F). Immunofluorescence analysis at 8-dpi showed that FAP^RFP^-WT transplants significantly reduced the number of embryonic myosin heavy chain (eMYHC) positive cells and increased CSAs in regenerating fibers compared to aged recipients transplanted with FAP^RFP^-KO (Figures 3I-3K). Consistently, recipients transplanted with FAP^RFP^-WT maintained significantly increased CSAs at 14-dpi when compared to FAP^REF^-KO transplants (Figures 3L and 3M). Finally, we compared the CSAs average and fiber size range results at 14-dpi to the original experiments without FAP-transplantation (i.e. CSA-means and fiber size range of data reported in Figure 1M). Aged mice receiving FAP^RFP^-KO transplants displayed significantly lower average CSA compared to young control mice and were at a similar level compared to aged TA without FAP transplant, whereas Periostin-producing FAP^RFP^-WT transplants significantly increased regenerating myofibers’ CSA compared to aged mice without transplanted FAPs or with FAP^RFP^-KO transplants (Figure 3N and S3G). Thus, reinstating Periostin as a niche factor ameliorates the age-related defect of myogenic regeneration, indicating that Periostin is a critical ECM-protein to orchestrate muscle regeneration.

### Periostin mediates immunomodulatory functions of the myogenic stem cell niche

Since regeneration is orchestrated by the interplay of multiple non-myofiber cell populations^47^, we performed droplet-based scRNA-seq of FACS-purified non-myofiber cells from 2.5- and 15-month-old female mice at 4-dpi, around the time when FAPs frequency peaks during muscle injury repair (Figures S4A and S4B). UMAP visualization, unsupervised Louvain-clustering and subsequent cell type annotation based on gene scoring of single cell transcriptomes resulted in the identification of distinct clusters representing the main nine, non-myofiber cell populations involved in muscle regeneration (Figures 4A, 4B, S4C and S4D; Tables S3 and S4) ^48^. Analysis of the cell-type proportions between age groups showed increased relative abundances of myeloid subsets, i.e. neutrophil (NP) and macrophage (MC) subsets in aged muscle, while lymphoid subpopulations, i.e. T-lymphocytes (TC) and natural killer cells (NK), showed lower relative frequencies (Figure 4C; Table S5). Considerable differences in gene expression patterns were observed among multiple cell types and when comparing the 2.5- and 15-months-old samples. For instance, analysis of enriched term annotations of downregulated DEGs from the FAP subset confirmed that aging negatively influences ECM regulation (Figure S4E, Table S6). As before, *Postn* was mostly expressed in FAPs and its expression was significantly reduced in aged FAPs (Figures 4D, 4E, and S4D; Table S4). To examine cell-cell interactions (CCI) between Periostin-secreting FAPs and other niche cells, we took advantage of a previously published murine atlas of age-related changes in intercellular communications, which uses an algorithm to infer CCI from published gene expression databases^49^. When selecting Periostin as a FAP-emitted ligand, high CCI-scores were identified for a potential autocrine FAP-FAP interaction and for paracrine FAP-endothelial cell (EC), FAP-MuSC and FAP-MC interactions mediated through different Integrin receptors (*Itga5*, *Itgβ1*, and heterodimers *Itgαv/β5*, *Itgαv/β*3) and the Epidermal growth factor receptor (*Egfr*; Figure 4F). Using aging as a regulatory factor of these interactions, statistical significance levels for each interaction showed a significant down-regulation of Periostin-based FAP-FAP, FAP-EC, FAP-MuSC and FAP-MC interactions, thus verifying our own experimental data on the auto- and paracrine roles of Periostin (Figure 4G). The age-dependent down-regulation of the interaction between FAPs and MCs through the predicted CCI between Periostin and the Itgαv/β5 heterodimer raised our interest as this was consistent with the alterations in immune subsets in our scRNA-seq data and offered the possibility of an immunomodulatory niche effect through the FAP-based release of Periostin (Figure 4G). We validated subset-specific expression patterns of all potential Periostin receptors in our scRNA-seq data and found that the proposed Itgαv/β5 heterodimer showed an expression pattern that would most closely reflect the effects of Periostin on the cell types that we also detected in our experimental analyses, i.e. FAPs, MuSCs, and MCs (Figure 4H).

**Figure 4:**
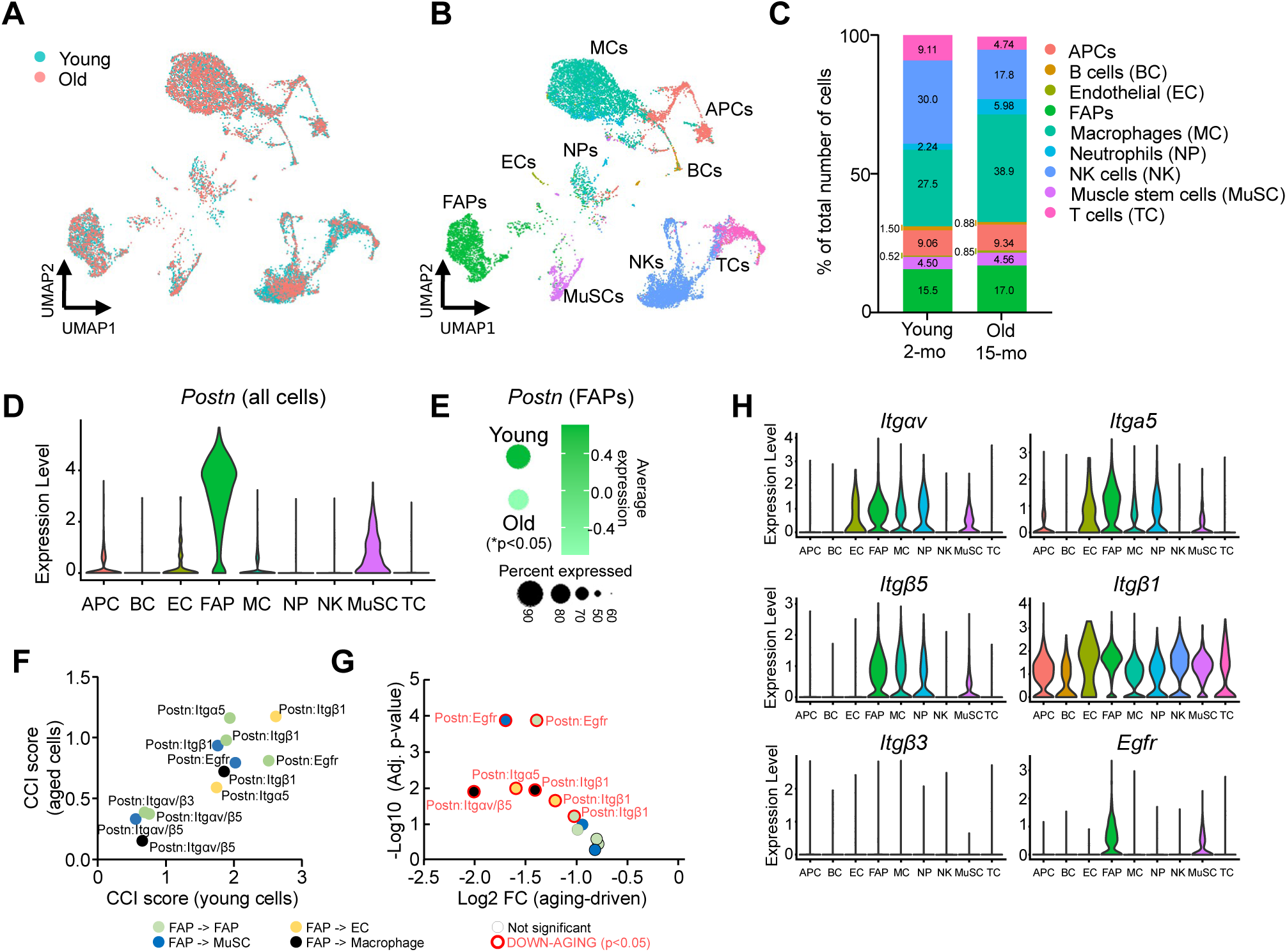
Computational analysis uncovers a Periostin-dependent FAPs-macrophage interaction. (**A**) UMAP of scRNA-seq data generated by droplet-based sequencing technology of all cells isolated from 4-dpi TA-muscle, integrating single cell transcriptomic data of young (2 months; blue-green) and aged (15 month; reds) female C57BL/6J mice. (**B, C**) Unsupervised Louvain clustering analysis and bar plot of population comparison in percentages of nine distinct cell populations identified in regenerating muscle. (**D**) Violin plot showing *Postn* expression levels across the nine cell populations. (**E**) Mean expression and percentage of cells expressing *Postn* in FAPs between 2- and 15-months groups (p-value: Wilcoxon rank sum test). (**F**, **G**) Age-dependent cell-cell interaction (CCI) scoring of ligand receptor interactions of Periostin with known receptors as a function of age (left panel, F) and interaction depiction of age-related changes in ligand-receptor-driven CCIs linking log fold changes and adjusted p-values (right panel, G) using a published input scRNA-seq dataset comparing young and aged female murine limb-muscle as described in^49^. Cell types within each CCI are color-indicated; red circles indicate statistically significant age-dependent changes in CCI scores. (**H**) Violin plot showing expression for *Itg*α*v*, *Itga5*, I*tgβ5*, *Itgβ1*, *Itgβ3*, and *Egfr* receptors across individual cells in the nine cell populations detected in our scRNA-seq dataset.

### Loss of Periostin impairs FAP-based immunomodulation by attenuating anti-inflammatory macrophages

Corroborating the FAP-MC interaction through Periostin, FACS-analyses of MCs populations confirmed the increase of MCs upon muscle injury in aged and also female *Postn^-/-^* mice compared to the respective control animals, while only trends or no differences were observed in male *Postn^-/-^* mice (Figures S5A-S5E). Next, we examined MC subsets representing functional changes from a pro-inflammatory (M1-like) profile to an anti-inflammatory (M2-like) phenotype during the course of muscle healing^25,50^. As expected, injury led to a rapid increase of CD45^+^CD11b^+^F4/80^+^ MCs, which over time shifted from a predominantly pro-inflammatory Ly6C^+^ population, which peaked at 1- and 3-dpi, to an anti-inflammatory Ly6C^-^ population which peaked at 4- and 8-dpi (Figure 5A). Importantly, aging as well as genetic ablation of *Postn* led to an attenuation of this shift in male and female mice, both of which resulted in significantly elevated levels of pro-inflammatory MCs at earlier time points and significantly lower levels of anti-inflammatory MCs at the later time points (Figures 5A-5C, S5F and S5G). We confirmed this delay in the formation of an anti-inflammatory, pro-regenerative MC profile by assessing CD206, a surface marker commonly found on anti-inflammatory MCs^51–53^. Aging and the deletion of *Postn* significantly reduced the accumulation of CD45^+^CD11b^+^F4/80^+^CD206^+^ MCs in the late, regenerative phase at 3-, 4- and 8-dpi, regardless of gender (Figures 5D-5F, S5H and S5I). Lastly, we examined MCs in our FAP transplantation model (Figure 3H) and detected no differences in overall MCs but a similar phenotype of significantly elevated CD206^+^ MCs in 8-dpi-injury sites of aged mice transplanted with *Postn*-expressing FAPs compared to *Postn*-deficient transplants (Figures 5G and 5H). These data reiterate the notion that loss of Periostin, similar to aging, delays the formation of a regeneration-supporting anti-inflammatory niche.

**Figure 5:**
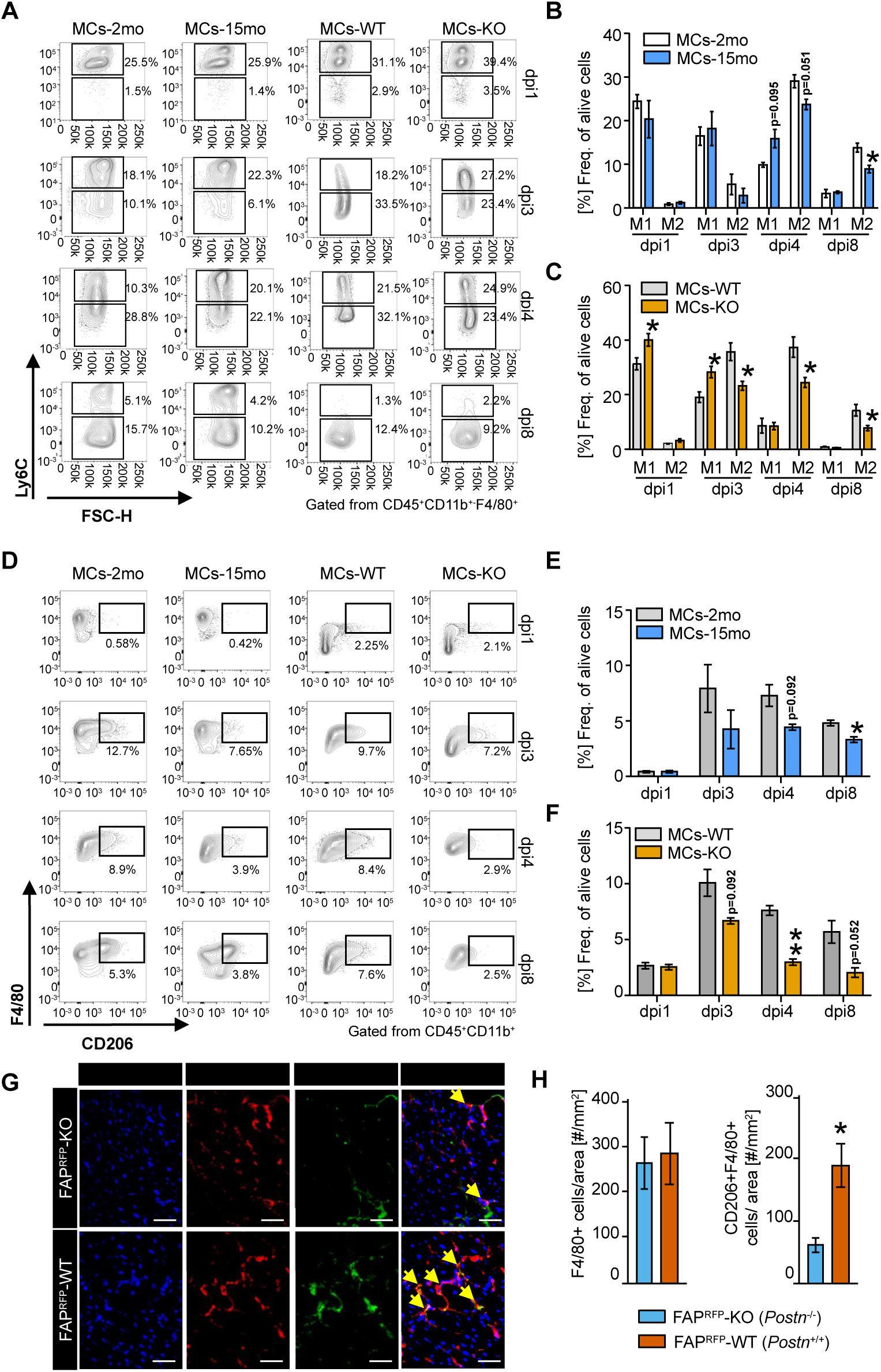
Aging and lack of Periostin impair macrophage polarization in regenerating muscle. (**A**) Representative density plots from FACs analyses of macrophage populations comparing either young (MCs-2mo) to aged (MCs-15mo), or young wildtype (MCs-WT) to littermate *Postn^-/-^* (MCs-KO) mice. Total MCs were initially gated as CD45^+^CD11b^+^F4/80^+^ and subsequently for Ly6C-expression to indicate Ly6C-high, pro-inflammatory (M1) vs. Lyc6-low, anti-inflammatory (M2) MCs at 1-, 3-, 4-, and 8-dpi. (**B**, **C**) Quantitative analysis of MC frequencies as described in panel A (n=3-4/group). Comparisons are young (MCs-2mo; white bars) to aged (MCs-15mo; blue bars) and wildtype (MCs-WT; gray bars) to Postn^-/-^ (MCs-KO; orange bars; also applies to panels E, F) mice. (**D**) Representative density plots of anti-inflammatory F4/80^+^CD206^+^ MCs comparing groups as described in panels A-C. MCs were initially gated as CD45^+^CD11b^+^. (**E**, **F**) Quantitative analysis of MCs frequencies as described in panel D (n=3-4/group). (**G**, **H**) Representative immunofluorescence images and quantifications of F4/80^+^ and F4/80^+^CD206^+^ MCs in injured site of 15-months old TA-muscle sections at 8-dpi transplanted with FAP^RFP^-WT (blue bar) or FAP^RFP^-KO (brown bar) at 1-dpi (red: F4/80, macrophage-marker; green: CD206, M2-marker; blue: DAPI, nuclei; n=4/group; yellow arrows indicate co-localization of F4/80^+^ and CD206^+^ MCs; scale bar: 20 µm). All animals were male, all data are presented as mean ± standard error of the mean (SEM), statistical significances are *p<0.05, **p<0.01, ***p<0.001.

### Periostin promotes anti-inflammatory macrophages through integrin-FAK-AKT mechano-signal transduction

To investigate the mechanism of Periostin-dependent control of macrophage polarization, we co-cultured wildtype and *Postn^-/-^*-FAPs with bone marrow-derived macrophages (BMDM) in a trans-well system and compared them to known pro- and anti-inflammatory stimuli. After 48h, FAPS-WT (*Postn^+/+^*), but not FAPs-KO (*Postn^-/-^*), significantly induced expression of anti-inflammatory MC markers *Mrc1* and *Cd163* while other anti-inflammatory genes, *Il10*, and several pro-inflammatory markers were not affected by the presence of FAPs (Figures 6A and S6A). Addition of species-specific recombinant Periostin proteins (mrPOSTN, hrPOSTN) to murine BMDM- and human, THP-1 cell line-derived macrophages resulted in a significant, dose-dependent increase in the expression of anti-inflammatory markers *Mrc1*, *Cd163* and *Il10*, but did not affect pro-inflammatory gene expression (Figures 6B, S6B and S6D). Integrins, and specifically Itgαv/β3, Itgαv/β5 heterodimers, have been linked to anti-inflammatory macrophage function^54,55^ and our own findings and published data indicate that Periostin as able to bind both integrin heterodimers^33,34^. Treatment of BMDM and THP-1 macrophages with Cilengitide, an inhibitor of ITGαv/β3 and ITGαv/β5 heterodimers^56,57^, resulted in a significant reduction of rPOSTN-induced expression of anti-inflammatory surface protein CD206 and similar effects on *Mrc1*, which encodes for CD206, and *Cd163* mRNA expression, whereas recombinant IL4-induced expression of these genes was not affected by Cilengitide and little or no effect on other pro-inflammatory genes was evident (Figures 6C-6E, S6C, S6E and S6F). Protein kinase B (AKT) and focal adhesion kinase (FAK) phosphorylation have been linked to anti-inflammatory macrophage polarization^58,59^ and both kinases are potential downstream targets of the Periostin-ITGαvβ3 and -αvβ5 axis^33,34,60^. Accordingly, rPOSTN acutely induced the phosphorylation of FAK at Tyr397 and of AKT at Ser437, which was reduced significantly after Cilengitide addition in BMDM and THP-1 macrophages (Figures 6F, 6G, S6G and S6H). Collectively, these results indicate that rPOSTN induces M2-like polarization through integrin-dependent FAK-AKT signaling in macrophages.

**Figure 6:**
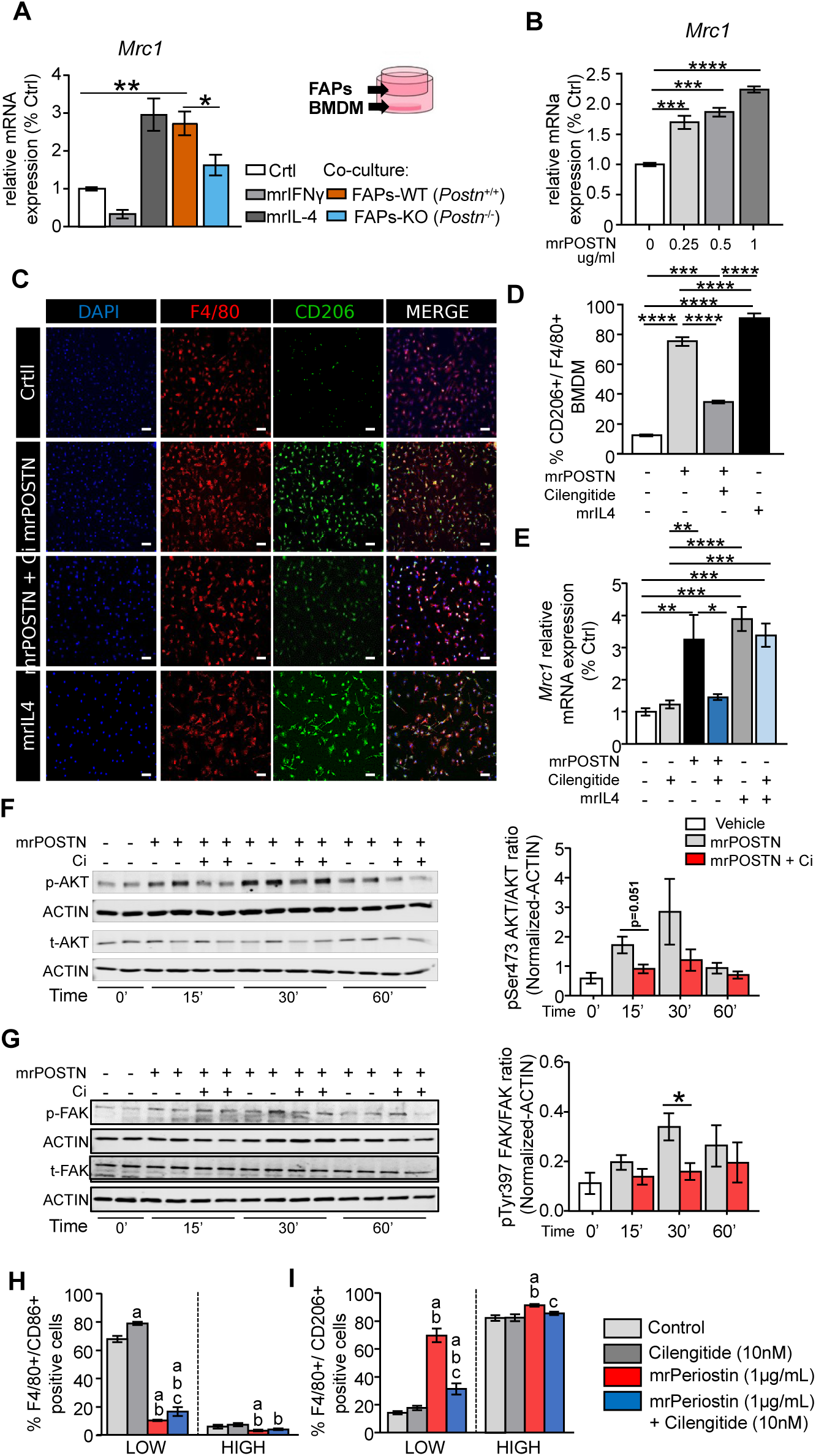
Periostin promotes anti-inflammatory niche transition through mechano-sensitive integrin-FAK-AKT signaling. (**A**) *Mrc1* (Mannose receptor c-type 1) mRNA expression in differentiated BMDM exposed to pro- and anti-inflammatory cytokines (mrIFNγ: 50 ng/mL; mrIL4: 10 ng/mL; n=8/treatment) or co-cultured with FAPs-WT or FAPs-KO (n=11, co-culture group) for 48h compared to untreated BMDM control (n=6) in 3 independent experiments. (**B**) *Mrc1* mRNA expression in differentiated BMDM treated with mrPOSTN at indicated concentrations for 48h (n=3/group). (**C**, **D**) Representative immunofluorescence images and quantification in untreated, differentiated BMDM control compared to a 48h treatment with mrPOSTN (1 µg/mL) alone or in combination with integrin receptor-inhibitor Cilengitide (10 nM). mrIL4 (10 ng/mL) was used as positive control of M2-polarization (green: CD206, M2-marker; red: F4/80, macrophage-marker; blue: DAPI, nuclei; n=3/treatment; scale bar: 50 µm). (**E**) *Mrc1* mRNA expression in untreated BMDM control compared to BMDM 48h-exposed to Cilengitide (10nM), mrPOSTN (1 µg/mL), mrPOSTN + Cilengitide (1 µg/mL + 10nM), mrIL4 (10 ng/ml) or mrIL4 + Cilengitide (10 ng/mL + 10 nM) (n=4/treatment). (**F, G**) Protein detection and quantification of phosphorylated Ser473-AKT (p-AKT) and total AKT (panel F) and phosphorylated Tyr397-FAK (p-FAK) and total FAK (panel G) in untreated compared to treated BMDM with mrPOSTN (1 µg/mL) alone or combined with Cilengitide (10 nM) for indicated times (Actin loading control; n=4; 2 independent experiments in panels F, G). (**H**, **I**) Immunofluorescence quantifications of pro-F4/80^+^CD86^+^ and anti-inflammatory F4/80^+^CD206^+^ MC polarization of differentiated BMDM encapsulated in soft or high stiffness hydrogels untreated (control) or treated with mrPOSTN (1 µg/mL) and Cilengitide (10 nM) alone or combined (mrPOSTN + Cilengitide) for 24h (n=12/group; 2 independent experiments). All data are presented as mean ± standard error of the mean (SEM), statistical significances are *p<0.05, **p<0.01, ***p<0.001, ****p<0.0001.

Aging and fibrotic scar tissue formation during muscle regeneration have been associated with altered muscle stiffness, somewhat resembling the reduced muscle stiffness we observed in our *Postn*^-/-^ mice (Figure S2F) ^12,23^. During muscle regeneration, a transient decrease in tissue stiffness at the injury site around 5-dpi, which subsequently rebounds to normal stiffness levels in fully regenerated muscle, has been reported recently^61^. To investigate the relationship of Periostin and the mechanical properties of the local niche, and their impact on macrophage polarization, BMDM were encapsulated in a three-dimensional ECM microenvironment using different alginate-based hydrogels with either low or high mechanical stiffness. Consistent with previous reports^62^, low stiffness hydrogels contained significantly fewer F4/80^+^/CD206^+^ anti-inflammatory MCs compared to high stiffness hydrogels, while the opposite was true for F4/80^+^/CD86^+^ pro-inflammatory MCs, which were more abundant under low stiffness and practically absent under high stiffness conditions. These data suggest that mechanical input from the microenvironment directly affects macrophage polarization and imply that loss of *Postn* may exert a similar effect in the muscle niche. Importantly, the addition of murine rPOSTN to the hydrogels resulted in a Cilengitide-sensitive, modulatory effect where pro-inflammatory polarization induced by low stiffness was significantly reduced. Instead, an anti-inflammatory MC profile was promoted under conditions of low stiffness, suggesting that Periostin acts as a niche factor that synchronizes biomechanical and immunomodulatory processes within the muscle niche (Figures 6H, 6I and S7A-S7C).

### Circulating Periostin may predict muscle dysfunction in geriatric patients

Circulating Periostin has been proposed as biomarker of several inflammatory, chronic and metabolic diseases^39–42^, but its role in relation to aging in mice and humans is only now emerging^22,43^. To explore the link between aging and muscle dysfunction, we measured circulating human Periostin (hPOSTN) in serum samples from two separate human study cohorts^63,64^. When comparing serum Periostin from young adults to healthy older, community-dwelling individuals, significantly lower levels of circulating hPOSTN were observed in aged individuals. Additionally, we found that increased physical activity in elderly individuals restored hPOSTN-concentrations to levels found in young adults (Figure 7A). As previous studies had reported elevated levels of circulating Periostin due to certain pathologies^65^, we also assessed serum hPOSTN in a group of healthy older individuals compared to age-matched individuals living in an acute geriatric hospital setting. No differences were found, indicating that circulating hPOSTN is not applicable as a generalized marker of impaired health status (Figure 7B). Next, we performed correlations including young adults to healthy older community-dwelling individuals only (from panel 7A), and we found an inverse correlation between age and circulating hPOSTN serum levels in women, but not in men, which could be due to lower inclusion numbers for men (Figure 7C). Using all individuals of this study cohort, we subsequently examined whether other metabolic and functional muscle parameters would also correlate with circulating hPOSTN. Indeed, hPOSTN correlated inversely with body mass index (BMI) and its serum concentration was positively corelated with skeletal muscle mass and grip strength in women, but not men (Figures 7D-7F). These data suggest that circulating Periostin levels could be used as a potential biomarker of skeletal muscle aging and the age-related sarcopenia syndrome.

**Figure 7:**
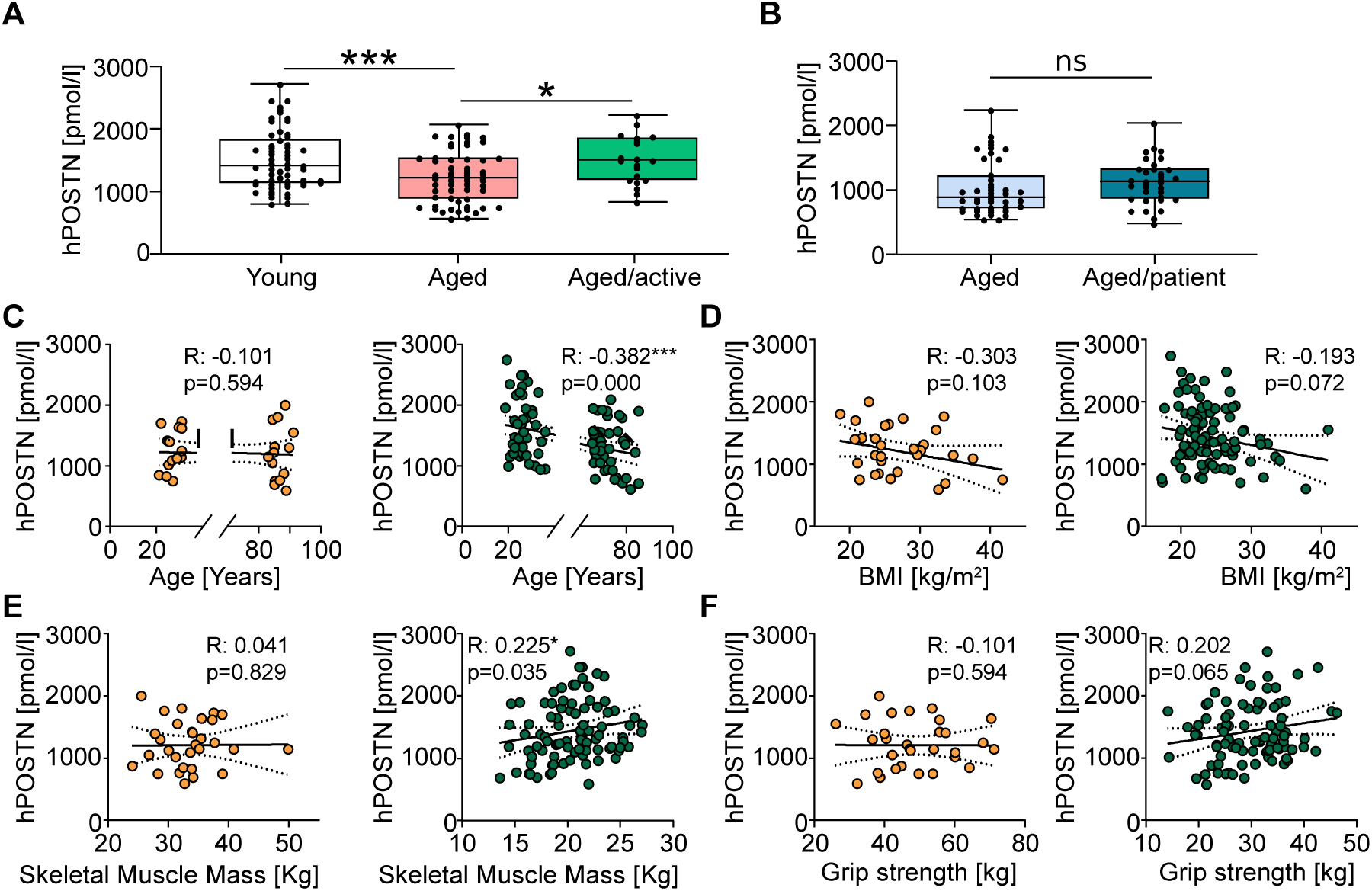
Circulating Periostin functions as biomarker of muscle dysfunction in elderly patients. (**A**) Serum levels of human Periostin (hPOSTN) in young adults (white; n=59; n=16 men; n=43 women), healthy aged, community-dwelling adults (red; n=59; n=14 men; n=45 women), and aged community-dwelling adults with high activity levels (Aged/active; green; n=20; n=5 men; n=15 women). (**B**) Serum hPOSTN levels in community-dwelling aged adults (light blue; n=44), and geriatric hospital patients (Aged/patient; dark blue; n=34). (**C-F)** Pearson correlations of serum hPOSTN levels and individuals’ age (panel C), body mass index (BMI, panel D), total skeletal muscle mass (panel E) and grip strength (panel F) separately in men (orange, left) and women (green, right), including young adults and aged community-dwelling individuals only, as described in panel A. All data are presented as mean ± standard error of the mean (SEM), statistical significances are *p<0.05, **p<0.01, ***p<0.001, ****p<0.0001.

## Discussion

Muscle injury repair requires complex interactions between different cell types residing within the stem cell niche^47^. For instance, extensive evidence exists for FAP-MuSC interactions to ensure successful myogenic regeneration^10,13–15,23,24,66^. Aging and other pathologies result in defective myogenesis due to stem cell niche dysfunction by inhibiting different aspects of myogenic stem cell maintenance, activation and differentiation^4^. We here report on a niche-dependent mechanism that integrates paracrine signals and mechano-signal transduction to impact on the levels of MuSCs and the myogenesis-regulatory FAPs, and by exerting immunomodulatory effects regulating the inflammatory profile of injury site-associated macrophages. De-synchronization of these processes creates a defective stem cell niche that phenocopies the detrimental effects of aging on myogenic regeneration, which result in enhanced fatty degeneration of muscle tissue instead.

Firstly, we show that loss of the matricellular protein, Periostin, which is secreted by muscle-resident FAPs, directly impinges on MuSCs, attenuating their ability to expand in response to injury and to differentiate into fully functional myofibers. These findings are consistent with recent studies showing that aging alters secretion of signaling molecules like the WNT1-inducible signaling pathway protein 1 (WISP1), bone morphogenetic protein 3 (BMP3b) and SPARC-related modular calcium-binding protein 2 (SMOC2) by FAPs in the stem cell niche, thereby reducing MuSCs’ regeneration capacity^10,11,67^.

Previous reports have established that aging and chronic disease prime FAPs to form adipocytes that accumulate between muscle fibers, leading to an impairment of muscle function^2,6,16^. In line with these observations, our animal models with loss of Periostin due to age or genetic targeting display defective autoregulation of FAPs, resulting in excessive FAP-expansion upon injury which feature an elevated commitment towards the adipogenic lineage at the expense of their ability to maintain appropriate myogenic regeneration capacity. This process results in the formation of a pro-fibro/adipogenic microenvironment with enhanced fibro/fatty infiltration instead of successful muscle repair. A tight regulation of FAP expansion and clearance is essential to maintain tissue integrity and prevent fibro/fatty degeneration. However, the mechanisms through which aging may affect FAPs’ expansive capacities in homeostatic and regenerating muscle tissues remain unclear. Previous reports have described a reduction of FAPs in uninjured muscle during aging^10,24^. Upon injury the situation could depend on the specifics of the experimental layout, physiological status and age, as age-related failure of FAPs to expand has been reported^10^, while our own observations suggest enhanced FAP expansion, which align with scRNA-seq and tissue-array data at steady-state in old muscle^8,68^. Intramuscular FAP transplants into mice with impaired myogenesis have been shown to rescue the normal characteristics of healthy muscle under homeostatic conditions and improved regeneration capacity^10,14,21^. Our FAP/MuSC co-cultures and FAPs-transplants into muscle injuries of aged mice help restore secretion of Periostin and are sufficient to reconstitute myogenesis and muscle regeneration. Our findings therefore support the concept that FAP-secreted ECM factors are necessary and sufficient to sustain myogenesis via the FAP-MuSC crosstalk.

Our work sheds additional light on the concept of ECM-linked factors that exert immunomodulatory effects by targeting the pro- to anti-inflammatory transition of macrophages within the muscle injury site. This process integrates mechanical properties of the ECM and its changes during muscle repair with mechanosensory signaling through AKT and FAK pathways. When disrupted, for instance through loss of Periostin, this results in a delay of the pro- to anti-inflammatory switch within the post-injury myogenic niche and is also observed in aging muscle. The crosstalk between FAPs and muscle-resident immune cell subsets to sustain adequate muscle regeneration has been reported on before^23,26–28^. Nevertheless, the exact mechanisms of interaction between FAPs and macrophage polarization associated with muscle regeneration capacity, and their alterations during the aging process, remain largely unknown. We here introduce the concept that niche factors like Periostin may help to synchronize the biological signaling processes of immunomodulation and biomechanics during muscle regeneration. Previous reports have shown that FAPs coordinate myeloid and lymphoid cell contributions to inflammatory processes during the regeneration process^23,26–28^. Our scRNA-seq analysis of regeneration in young and aged muscle shows that myeloid and lymphoid subsets are the main clusters affected by aging. In line with these observations, experimental reconstitution of macrophage-derived signaling molecules growth differentiation factor 3 (GDF3) and mesencephalic astrocyte-derived neurotrophic factor (MANF) improved regeneration^31,69^. These results also support the concept of reinstating a normal macrophage function to ensure successful regeneration in the context of aging. This could be achieved by targeting regulatory FAPs to provide appropriate immunomodulatory signals, such as Periostin, to orchestrate the pro- to anti-inflammatory macrophage transition. In our hands, FAPs transplantation with functional Periostin expression into aged, post-injury muscle restored a timely onset of anti-inflammatory niche formation and successful regeneration. These findings are in line with previous results showing that Periostin could support a shift from a pro-towards an anti-inflammatory phenotype to sustain heart, kidney and bladder tissue regeneration, although a molecular mechanism has so far not been reported^51–53^. Thus, our data link FAPs-derived Periostin secretion to the muscle stem cell niche with synchronizing effects on FAPs, MuSCs and macrophages during muscle healing. Periostin exerts its function through integrin receptors and mainly through αvβ3 and αvβ5 heterodimers in different biological contexts^33–35^. Analyses of CCIs in the muscle stem cell niche corroborate that Periostin is a FAPs-secreted ligand that mediates intercellular communication between FAPs, MuSCs and macrophages. Interestingly, the Periostin:Itgαv/β5 interaction, which mediates the crosstalk between FAPs and macrophages, declines with aging in the model, thus mirroring our own experimental evidence and presenting a new mechanism of age-related macrophage dysfunction and niche deterioration where a relative lack of Periostin fails to activate FAK-AKT signaling. This in turn attenuates the transition from a pro-towards an anti-inflammatory milieu in the regenerating muscle niche and is in line with other studies that implicate Periostin in regulation of macrophage function in other organs and physiological contexts^34,51,59^. Corresponding immunomodulatory mechanisms have been described where the integrin-αvβ5 heterodimer regulates anti-inflammatory macrophages via a PPAR-γ dependent mechanism^55^, suggesting that this signaling pathway may also integrate different immunomodulatory signals during myogenic regeneration.

In addition to its role as a signaling molecule, our data also suggest a link between Periostin and mechanosensory signal transduction. The ability of matrices with different mechanical properties to regulate macrophage polarization is well-documented in the literature^62^. Molecules like Periostin could offer an additional layer of regulatory input that offsets or enhances the role of the local ECMs’ mechanical profile on inflammatory processes, including macrophage functions, thereby also impacting on myogenic stem cells and fatty degeneration of muscle. Aging increases muscle stiffness^12^, a mechanical property which would normally promote anti-inflammatory macrophages. It should be noted that such a situation would only occur in aged muscle at steady-state, i.e. non-injury conditions, and has indeed been reported^70^. In intact muscle, however, overall numbers of tissue macrophages are comparably low. Conversely, post-injury muscles experience dynamic ECM-remodeling, marked by an initial, but transient decline of stiffness at 5-dpi^61^, which in fact coincides with the pro- to anti-inflammatory niche/ macrophage transition. As low stiffness would preferentially stimulate pro-inflammatory macrophages, the presence of Periostin could mediate or at last accelerate the transition to an anti-inflammatory microenvironment.

Lastly, circulating Periostin has been proposed as a potential biomarker in some instances such as for cardiovascular risk, early cancer detection, or different type 2 inflammatory diseases^39–42^. Our own and other previous work suggests that reduced gene expression levels of *Postn* are reliable predictors of aging and associated metabolic disorders^22,43^. Thus, it stands to reason that loss of circulating Periostin may also inform on the progression of muscle degeneration as a result of age-related sarcopenia. It should be noted that increased levels of Periostin have also been proposed as a biomarker of disease progression in older adults, including cognitive decline^65^, suggesting that associations such as those found in our cohorts need to be evaluated in more detail. In conclusion, our study presents a novel role of FAP-based synchronization of the muscle stem cell niche, which links impaired regeneration to age-related niche deterioration and fatty degeneration of skeletal muscle. Our results provide mechanistic insights into a Periostin-dependent crosstalk between FAPs and macrophages that is impaired during aging, which integrates paracrine niche effects and mechano-sensing to orchestrate the precisely timed steps of successful myogenic regeneration instead of fibro-fatty infiltration.

## Acknowledgments

The authors thank A. Kretschmer, S. Marg, S. Richter, A. Weber and N. Dittberner for expert technical assistance. This work was supported by grants from the German Research Foundation (DFG) as individual grants and within the Collaborative Research Centre CRC1444 (project IDs 323196138, 249509554, and 427826188), and by a grant from the Leibniz Association (ID K398/2021, to TJS). Grants within the German Center for Diabetes Research (DZD) funded by the German Ministry of Education and Research (BMBF) and the State of Brandenburg (DZD grant IDs 82DZD00302, 82DZD03D03 and 82DZD03C3G) are acknowledged.

## Author Contributions

F.G.-C. and T.J.S. conceived the study and wrote the manuscript. F.G.-C. conducted the majority of experiments. S.G., G.L.-G., A.-M.J., C.G., S.P.-K., A.T., A.G., G.K. and M.Os., contributed research data to this study. G.S., L.V. und V.B. performed scRNA-seq experiments. M.L. and M.Ov. performed computational analyses of scRNA-seq data.

C.H., U.M.-W., K.N., provided human serum samples and helped with data analysis. S.K., A.S., S.S., K.S.-B., A.C., R.S.K., and G.N.D. contributed valuable materials and expertise to the article.

