## Supplementary Information: Figures, legends, materials and methods for "Aging impairs skeletal muscle regeneration by promoting fibro/fatty degeneration and inhibiting inflammation resolution via fibro-adipogenic progenitors"

### Supplementary figure 1

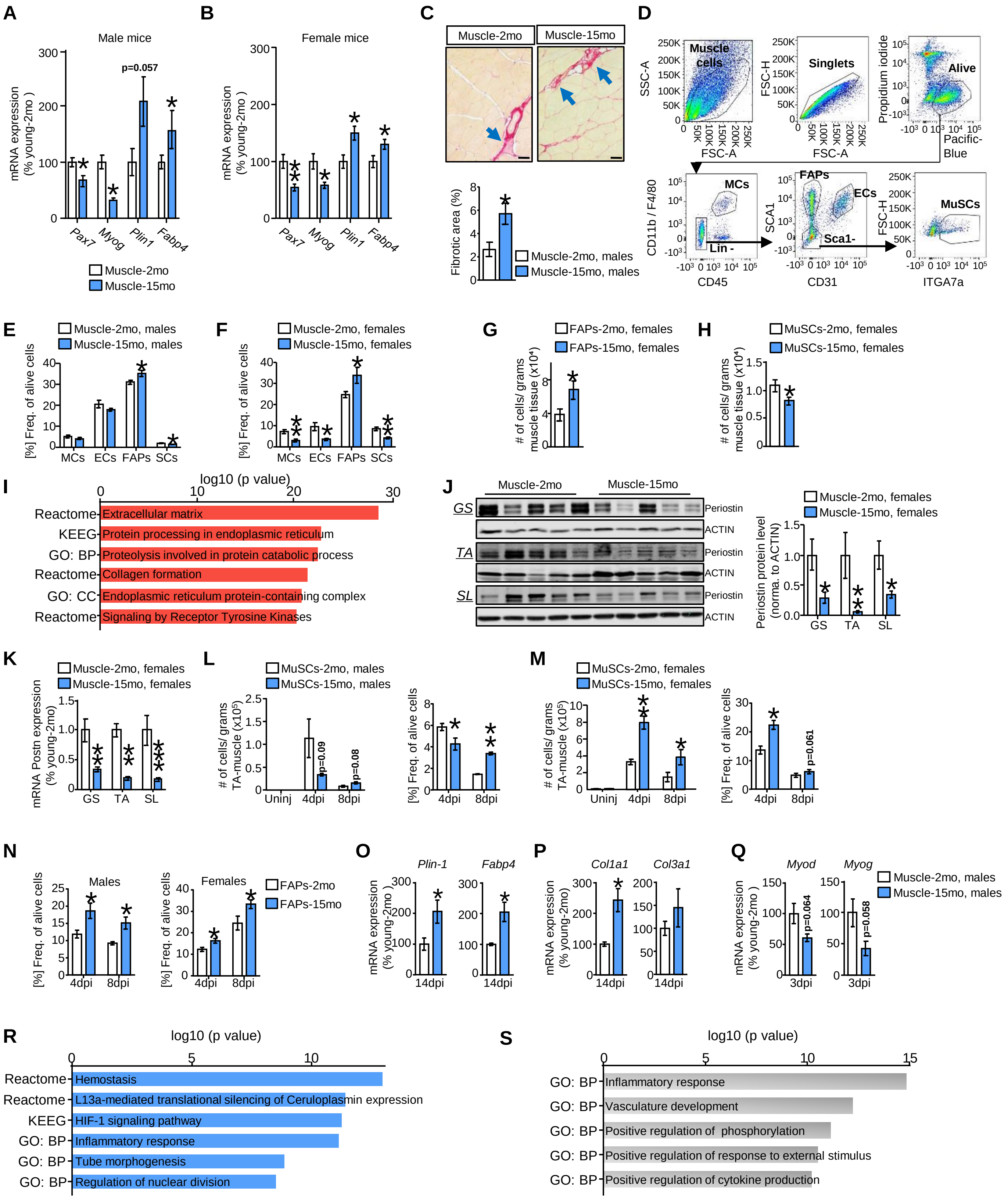

### Supplementary figure 2

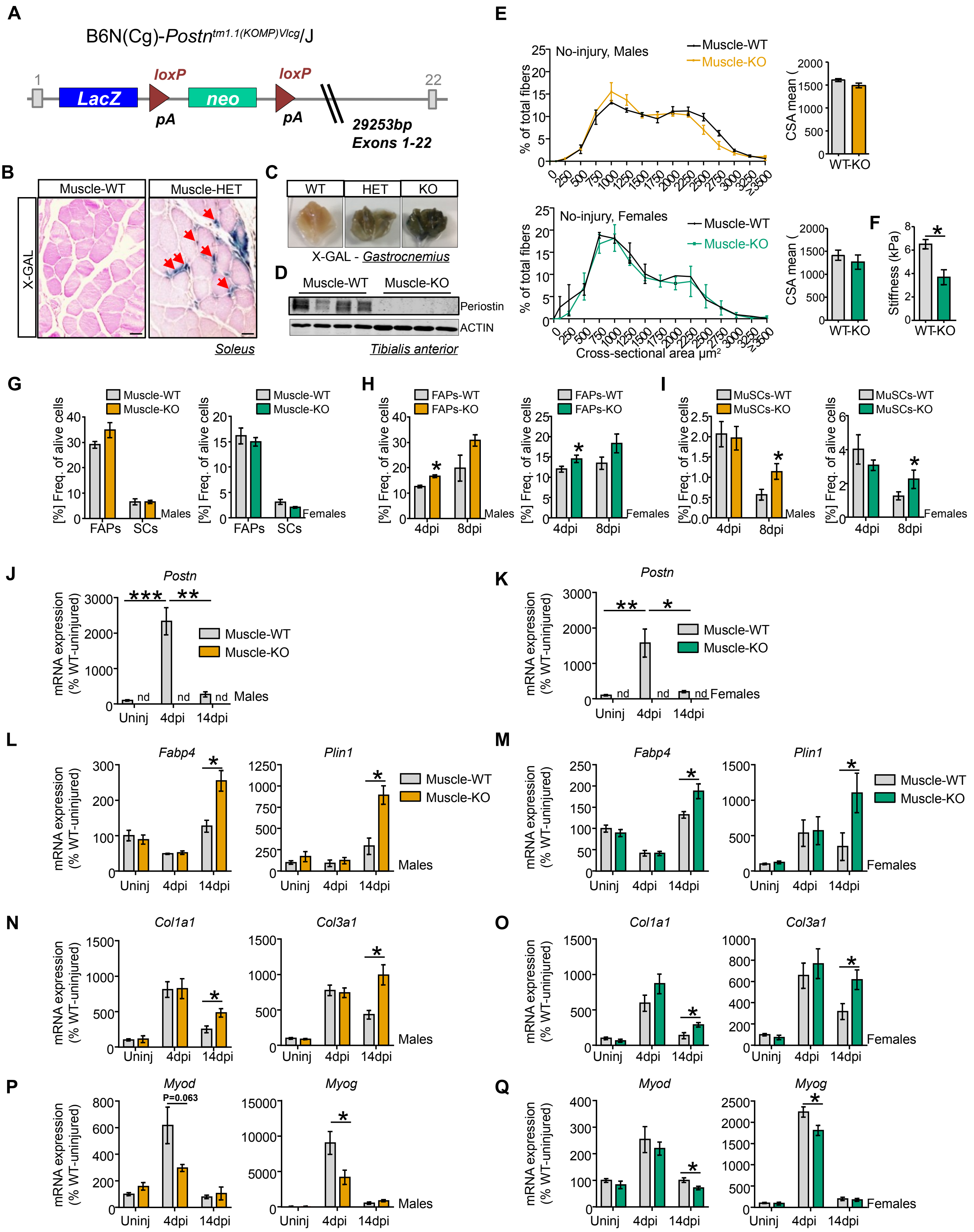

### Supplementary figure 3

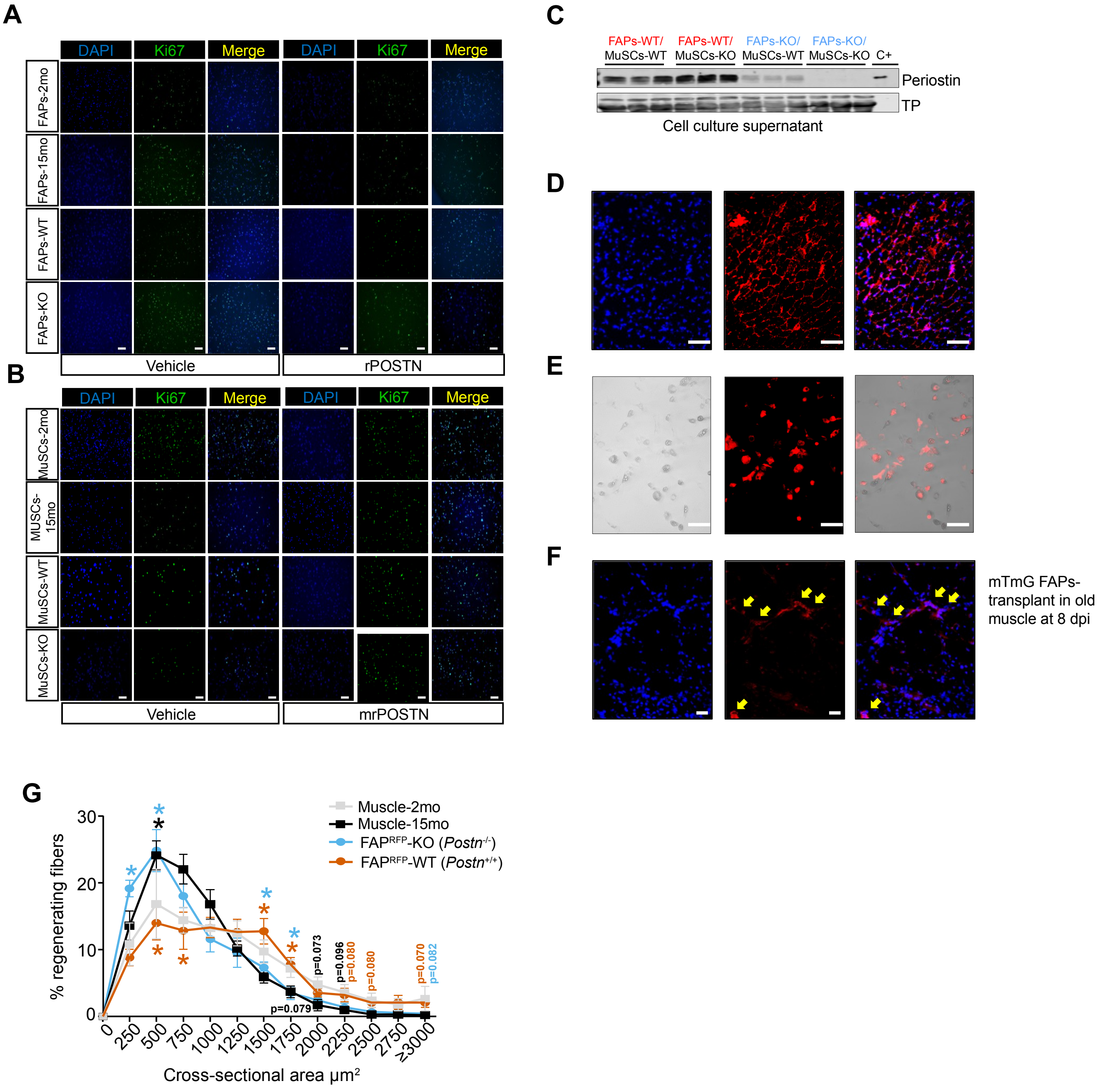

**Supplementary Figure 4**

**A**

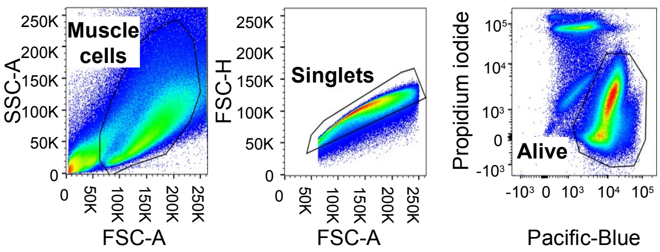


**B**

| Cell type and condition | Estimated Number of Cells | Mean Reads per Cell | Median Genes per Cell | Reads Mapped Uniquely to Genome | Median UMI Counts per Cell |
|---|---|---|---|---|---|
| Total alive cells in muscle - Young | 9,202 | 26,135 | 2,024 | 75.9% | 5,858 |
| Total alive cells in muscle - Old | 7,209 | 36,420 | 2,493 | 75.5% | 8,416 |

**C**

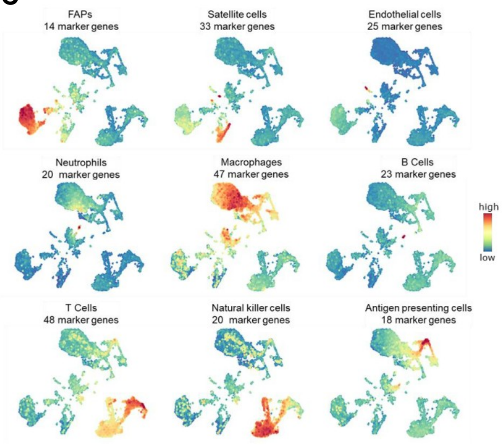


**D**

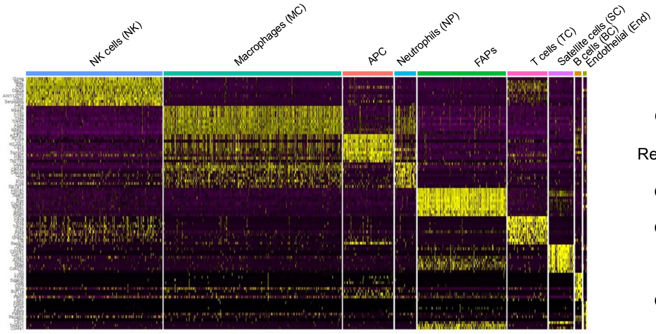


**E**

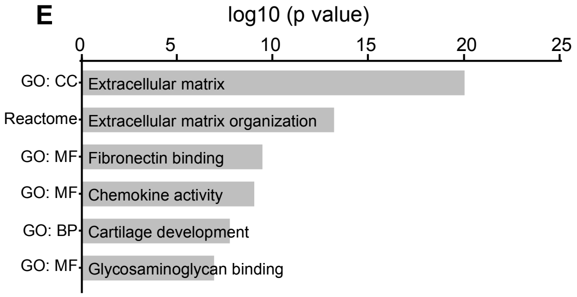

#### Supplementary figure 5

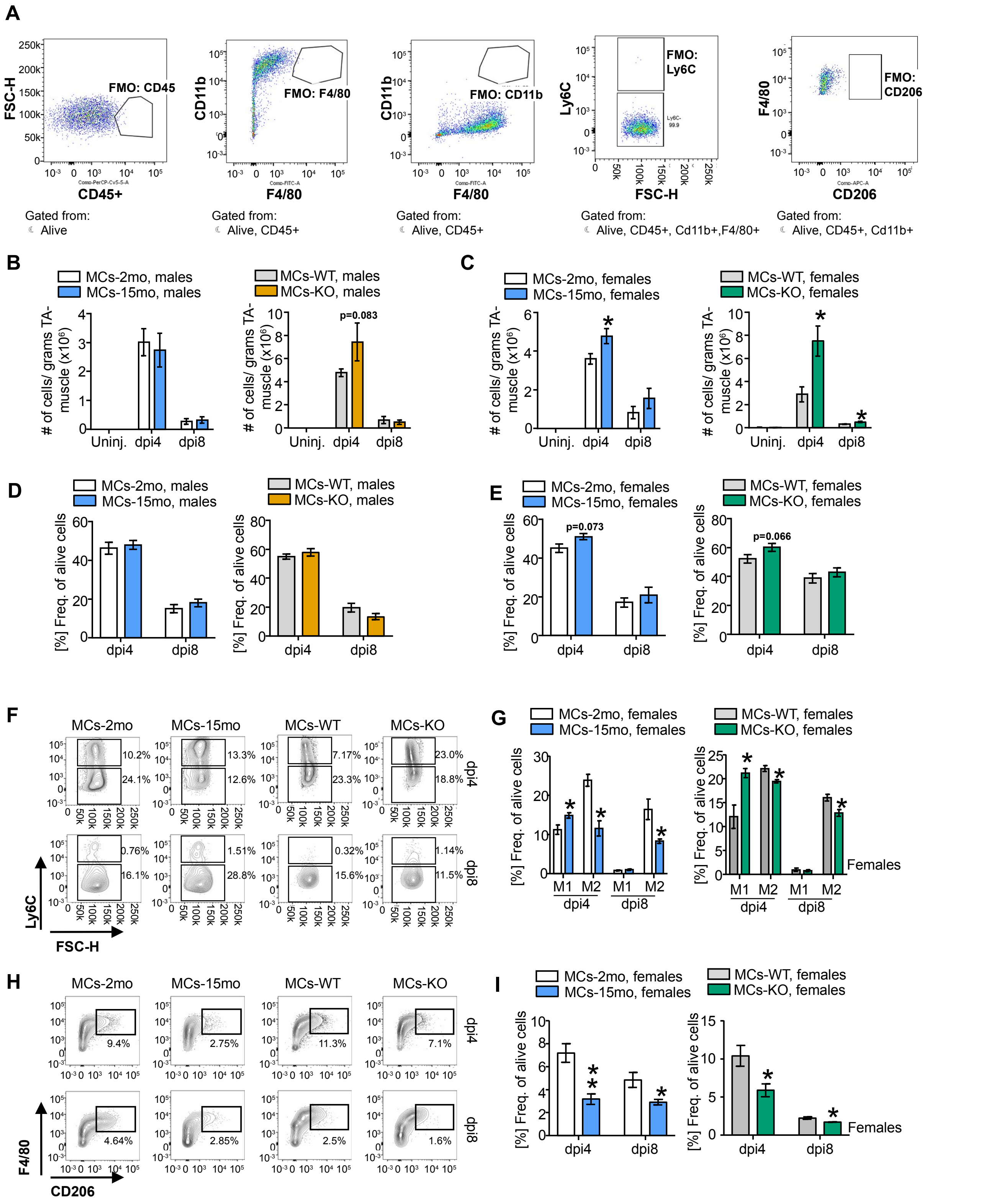

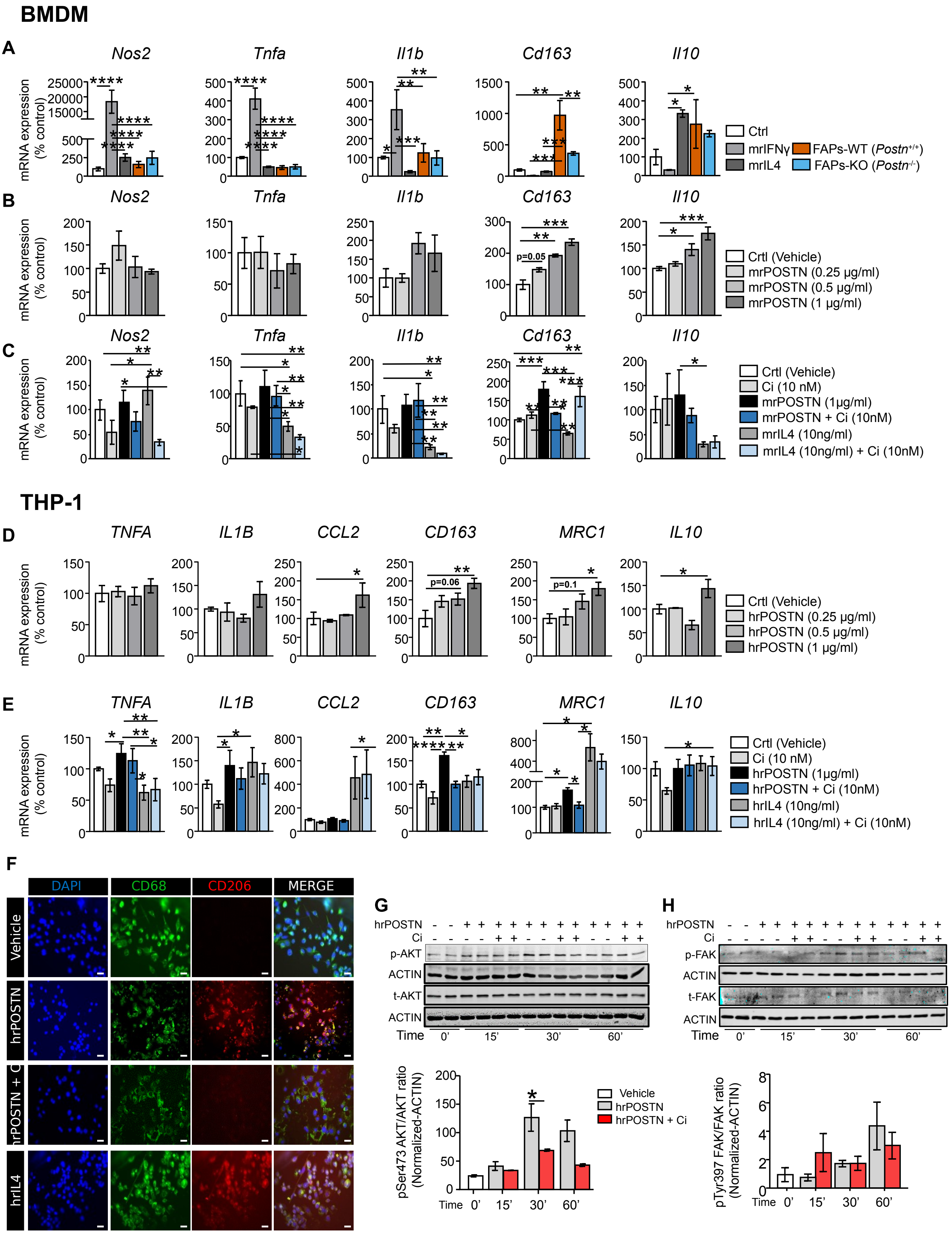
BMDM
A
Nos2
Tnfa
Il1b
Cd163
Il10
mRNA expression (% control)
Ctrl
mrIFNγ
mrIL4
FAPs-WT (Postn+/+)
FAPs-KO (Postn-/-)
B
Crtl (Vehicle)
mrPOSTN (0.25 µg/ml)
mrPOSTN (0.5 µg/ml)
mrPOSTN (1 µg/ml)
p=0.05
C
Crtl (Vehicle)
Ci (10 nM)
mrPOSTN (1µg/ml)
mrPOSTN + Ci (10nM)
mrIL4 (10ng/ml)
mrIL4 (10ng/ml) + Ci (10nM)
THP-1
D
TNFA
IL1B
CCL2
CD163
MRC1
IL10
Crtl (Vehicle)
hrPOSTN (0.25 µg/ml)
hrPOSTN (0.5 µg/ml)
hrPOSTN (1 µg/ml)
p=0.06
p=0.1
E
Crtl (Vehicle)
Ci (10 nM)
hrPOSTN (1µg/ml)
hrPOSTN + Ci (10nM)
hrIL4 (10ng/ml)
hrIL4 (10ng/ml) + Ci (10nM)
F
DAPI
CD68
CD206
MERGE
Vehicle
hrPOSTN
hrPOSTN + Ci
hrIL4
G
hrPOSTN
Ci
p-AKT
ACTIN
t-AKT
ACTIN
Time
0'
15'
30'
60'
pSer473 AKT/AKT ratio (Normalized-ACTIN)
Vehicle
hrPOSTN
hrPOSTN + Ci
H
p-FAK
t-FAK
pTyr397 FAK/FAK ratio (Normalized-ACTIN)

### Supplementary figure 7

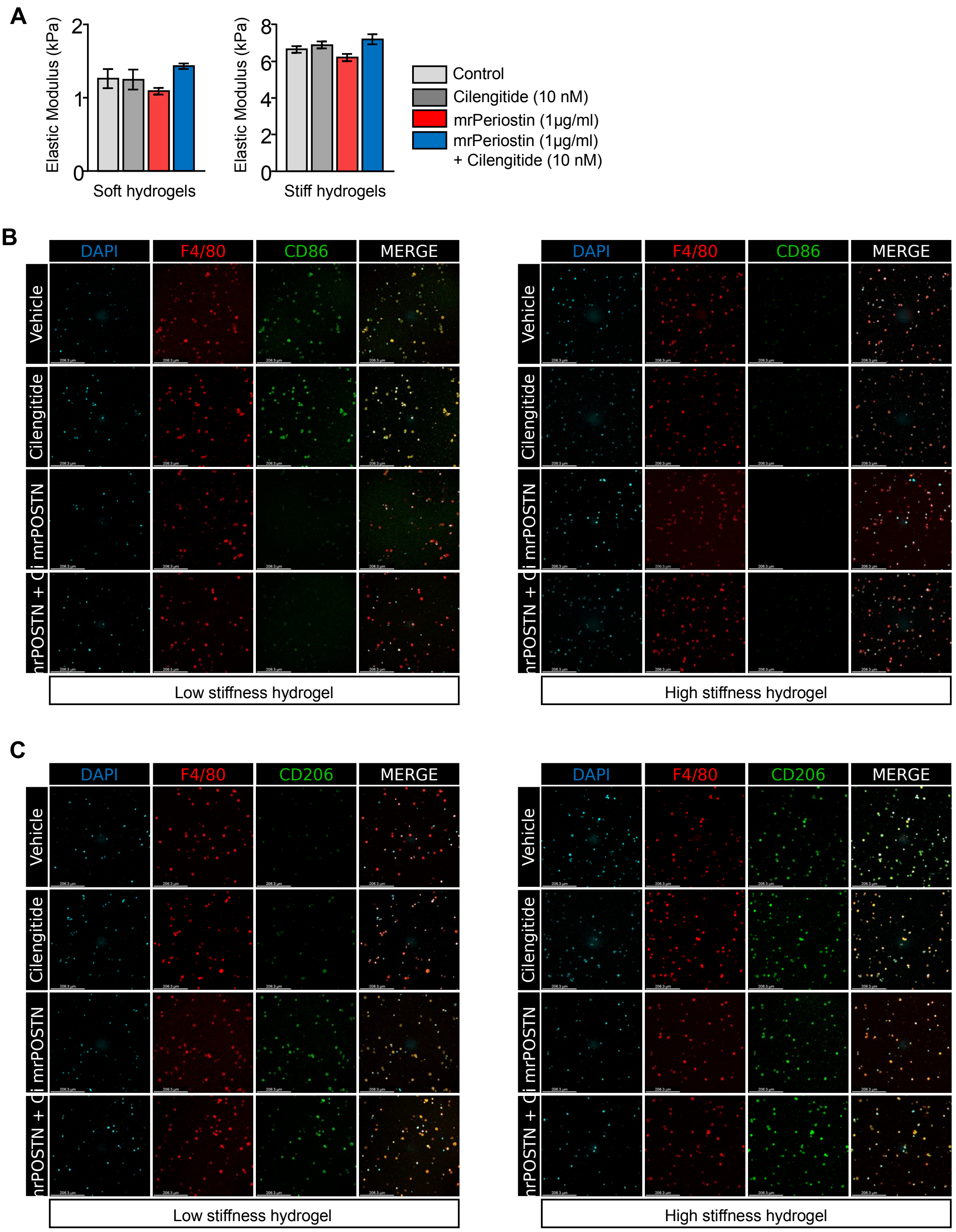

**Supplementary Materials Legends**

**Supplementary Figures**

**Supplementary Figure 1** (r**elated to Main Figure 1)**

(**A**, **B**) mRNA expression of myogenic markers *Pax7* (Paired box 7) and *Myog* (Myogenin) (upper panel) and adipocyte markers *Plin1* (Perilipin 1) and *Fabp4* (Fatty acid binding protein 4) (bottom panel) in uninjured gastrocnemius muscle samples of young (2 months, white bars) and aged (15 months, blue bars) male (panel A, n=6/group) and female mice (panel B; n=6/group).

(**C**) Representative Sirius red staining images (upper panel) and fibrotic depositions quantification (bottom panel) in gastrocnemius muscle of young (2 months) and aged (15 months) male mice (blue arrows indicate fibrosis accumulation; n=3/group; scale bar: 100 µm).

(**D**) Flow cytometric separation strategy for $CD45^+CD11b^+F4/80^+$ macrophages (MCs), $CD45^-CD31^-SCA1^+$ fibro adipogenic progenitor cells (FAPs), $CD45^-SCA1^-CD31^+$ endothelial cell (ECs), and $CD45^-CD31^-SCA1^-ITGA7^+$ muscle-myogenic stem cells (MuSCs) with typical prior gating of intact muscle cells, singlets and alive cells based on pacific blue (positive) and propidium iodide (negative) staining.

(**E**, **F**) FACS-analysis of frequencies of MCs, ECs, FAPS, and MuSCs as fraction of all alive cells in uninjured hindlimb muscles from young and aged male and female mice. (male: n=14/group; female: n=8/group).

(**G**, **H**) Flow cytometric analysis of the $CD45^-CD31^-SCA1^+$ FAPs and $CD45^-CD31^-SCA1^-$ $ITGA7^+$ MuSCs cells per grams of uninjured hindlimb muscles from young and aged female mice (n=8/group).

(**I**) Enriched terms and pathways of downregulated genes (fold change ≤ -1.5; p-value ≤ 0.05) in microarray analysis of freshly sorted FAPs of young (6 weeks) and aged (15 months) mice isolated from uninjured hindlimb muscles. Gene Ontology (GO) terms of biological process (BP) and cellular component (CC), pathway enrichment of Kyoto Encyclopedia of Genes and Genomes (KEEG) and reactome database using Metascape gene annotation and analysis tools.

(**J**) Periostin protein detection by western blot and quantification in uninjured gastrocnemius (GS), tibialis anterior (TA) and soleus (SL) muscles of young and aged female mice. Actin was used as loading control (n=5/group).

(**K**) *Postn* mRNA expression in uninjured gastrocnemius (GS), tibialis anterior (TA) and soleus (SL) muscles of young and aged female mice (n=5/group).

(**L**) Flow cytometric analysis of CD45$^-$CD31$^-$SCA1$^-$ ITGA7$^+$ MuSCs frequencies in uninjured (uninj) or at 4- and 8-dpi TA-muscle of young (2 months) and aged (15 months) male mice expressed per grams of TA-muscle weight (left panel) or as frequency of total alive cells (right panel) (n=3/group for uninjured [only in per tissue weight quantification] and 8-dpi; n=6/group for 4-dpi).

(**M**) Flow cytometric analysis of CD45$^-$CD31$^-$SCA1$^-$ ITGA7$^+$ MuSCs frequencies in uninjured (uninj) or at 4- and 8-dpi TA-muscle of young (2 months) and aged (15 months) female mice expressed per grams of TA-muscle weight (left panel) or as frequency of total alive cells (right panel) (n=4/group for uninjured [only in per tissue weight quantification], 4- and 8-dpi).

(**N**) Flow cytometric analysis of CD45$^-$CD31$^-$SCA1$^+$ FAP frequencies in 4- and 8-dpi TA-muscle of young (2 months) and aged (15 months) male (left panel; n=6/group 4-dpi, n=3/group 8-dpi) and female (right panel; n=4/group for 4- and 8-dpi) mice expressed as frequency of total alive cells.

(**O**, **P**, **Q**) mRNA expression of adipogenic marker genes *Plin1* (Perilipin 1) and *Fabp4* (Fatty acid binding protein 4) (panel O), fibrosis markers genes *Col1a1* (Collagen type I alpha 1 chain) and *Col3a1* (Collagen type III alpha 1 chain) (panel P) in 14-dpi TA-muscle and myogenic marker genes *Myog* (Myogenin) and *Myod* (Myogenic differentiation 1) (panel Q) in 3-dpi TA-muscle from young and aged male mice (n=4/group in 3- and 14-dpi).

(**R**, **S**) Enriched terms and pathways of upregulated genes (fold change ≥ 1.5, p value ≤ 0.05) and secretome annotation (panel S) from the scRNA-seq dataset of FACS-purified FAPs from young compared to aged male mice at 4-dpi. Gene Ontology terms of biological process (BP), pathway enrichment of Kyoto Encyclopedia of Genes and Genomes (KEEG) and reactome database using Metascape gene annotation and analysis tools.

All animals for experiments were young (2 months) and old (15 months) mice, unless otherwise indicated. All data are presented as mean ± standard error of the mean (SEM), statistical significances are *p<0.05, **p<0.01, ***p<0.001.

**Supplementary Figure 2** (r**elated to Main Figure 2)**

(**A**) Allele Details of *Postn* gene ablation by insertion of a LacZ reporter expression cassette to generate the transgenic mouse strain B6N(Cg)-Postn$^{tm1.1(KOMP)Vlcg}$/J with *Postn* gene inactivation and expression of LacZ-encoded beta-galactosidase under control of the *Postn* gene promoter.

(**B**) Representative X-Gal staining and hematoxylin/eosin counterstain of soleus muscle tissue cross-section collected from heterozygous *Postn*-knockout allele carriers (Muscle-HET) and wildtype littermates (Muscle-WT). Red arrows indicate X-Gal$^+$ areas with typical blue staining evident only in allele carriers.

(**C**) Whole-mount X-Gal staining of gastrocnemius muscles of wildtype controls (WT) as well as heterozygous (HET; *Postn*$^{+/-}$) and homozygous (KO; *Postn*$^{-/-}$) mutant allele carriers.

(**D**) Periostin protein detection by western blot in tibialis anterior muscle of female wildtype (WT) and *Postn*$^{-/-}$ (KO) mice, using Actin as loading control (n=4/group).

(**E**) CSA quantification of fiber size ranges and CSA averages in tissue sections of uninjured TA-muscle from littermate wildtype control mice of both genders (gray bars, black lines) in comparison to male (orange bar/line) and female (green bar/line) *Postn*$^{-/-}$ mice (n=3/group).

(**F**) Quantification of the TA-muscle stiffness calculated from assessment of elastic modulus after mechanical load in female wildtype controls (WT, gray bar) and *Postn*$^{-/-}$ (KO, green bar) female mice (n=3/group).

(**G**) Flow cytometric analysis of CD45$^{-}$CD31$^{-}$SCA1$^{+}$ FAPs and CD45$^{-}$CD31$^{-}$SCA1$^{-}$ ITGA7$^{+}$ MuSCs frequencies in uninjured hindlimb muscles from male (orange bars) and female (green bars) *Postn*$^{-/-}$ (KO) mice and corresponding wildtype (WT) littermates (gray bars) (males: n=3/group; females: n=3-5/group).

(**H**) Flow cytometric analysis of CD45$^{-}$CD31$^{-}$SCA1$^{+}$ FAP frequencies expressed as percentages of total alive cells in injured TA-muscle at 4- and 8-dpi in male (orange bars) and female (green bars) *Postn*$^{-/-}$ (KO) mice and their corresponding wildtype (WT) littermates (gray bars) (males: n=3-4/group; females: 4-dpi n=10/group, 8-dpi n=4/group).

(**I**) Flow cytometric analysis of CD45$^{-}$CD31$^{-}$SCA1$^{-}$ ITGA7$^{+}$ MuSCs cell frequencies expressed as percentages of total alive cells in injured TA-muscle at 4- and 8-dpi in male (orange bars) and female (green bars) *Postn*$^{-/-}$ (KO) mice and their corresponding wildtype (WT) littermates (gray bars) (males: n=3-4/group; females: 4-dpi n=10/group, 8-dpi n=4/group).

(**J**, **K**) *Postn* mRNA expression in uninjured (uninj) TA-muscle and at 4- and 14-dpi in male (orange bars) and female (green bars) *Postn*$^{-/-}$ (KO) mice and their corresponding wildtype (WT) littermates (gray bars) (males: uninj, 4-dpi n=4/group, 14-dpi n=3; females: uninj, 4-dpi n=7/group, 14-dpi n=8/group; nd: no expression detected).

(**L**-**Q**) mRNA expression of adipogenic marker genes *Plin1* (Perilipin 1) and *Fabp4* (Fatty acid binding protein 4) (panel L, males; panel M, females), fibrosis marker genes *Col1a1* (Collagen type I alpha 1 chain) and *Col3a1* (Collagen type III alpha 1 chain) (panel N, males; panel O, females), and myogenic marker genes *Myog* (Myogenin) and *Myod* (Myogenic differentiation 1) (panel P, males; panel Q, females) in uninjured (uninj) TA-muscle and at 4- and 14-dpi in in male (orange bars) and female (green bars) *Postn*$^{-/-}$(KO) mice and their corresponding wildtype (WT) littermates (gray bars) (males: uninj, 4-dpi n=4/group, 14-dpi n=3/group; females: uninj, 4-dpi n=7/group, 14-dpi n=8/group).

All animals for experiments were between 2 and 3 months of age, unless otherwise indicated. All data are presented as mean ± standard error of the mean (SEM), statistical significances are *p<0.05, **p<0.01, ***p<0.001.

**Supplementary Figure 3** (r**elated to Main Figure 3)**

(**A**, **B**) Representative immunofluorescence images of FAPs and MuSCs that were FACS-purified from either young (2 months) and aged (15 months) mice (upper panel group) or comparing Periostin wildtype (WT) and *Postn*$^{-/-}$ (KO) (bottom panel group) mice and cultured for 48h in growth medium with solvent control (vehicle) or with the addition of murine recombinant (mr)POSTN (0.5 µg/ml; green: Ki67, proliferation marker; blue: DAPI, nuclei; scale bar: 200 µm).

(**C**) Periostin protein quantification by western blot in supernatants of FAPs-WT or FAPs-KO co-cultured with MuSCs-WT or MuSCs-KO in myogenic differentiation medium for five days. Supernatants were collected 48h after last medium change (representing days 3-5 of differentiation) (C+ indicates mrPOSTN loading as positive control; TP: total protein staining as loading control).

(**D**) Representative immunofluorescence images of TA-muscle sections from the mTmG transgenic mouse model constitutively expressing the membrane-bound red fluorescent protein, tdTomato (red: tdTomato fluorescence; blue: DAPI, nuclei; scale bar: 100µm).

(**E**) Representative images of cultured FAPs isolated form mTmG mice displaying spontaneous differentiation to adipocytes in growth medium in light microscopy (left panel), red epifluorescence from tdTomato transgene (middle panel) and merge (right panel; scale bar: 100µm).

(**F**) Representative immunofluorescence images of TA-muscle sections from 15 months-old mice transplanted with FAP$^{RFP}$-WT (Main Figure 3) at 8-dpi. Yellow arrows indicate *in situ* presence of transplanted tdTomato$^{+}$ FAPs between regenerating myofibers at injury site (red: tdTomato detection; blue: DAPI, nuclei; scale bar: 20 µm).

(**G**) Combined comparisons of CSA quantification of fiber size ranges in injury sites at 14-dpi of data shown in Figure 1M (TA-muscle sections from 2- and 15-months old mice) and Figure 3M (TA-muscle sections from 15-months old mice transplanted with FAP$^{RFP}$-WT or FAP$^{RFP}$-KO). Data shown combine CSAs of 2-months TA muscle (grey square/line; n=4), 15-months TA muscle (black square/line; n=4), 15-months TA muscle transplanted with FAP$^{RFP}$-KO (blue circle/line; n=5), 15-months TA muscle transplanted with FAP$^{RFP}$-WT (orange circle/line; n=6). Data are presented as mean ± standard error of the mean (SEM), statistical significances are *p<0.05. Black color (*) represents statistical significances between TA muscle aged 2- vs. 15-months; orange color (*) represents statistical significances between 15-months vs. 15-months muscle transplanted with $FAP^{RFP}$-WT; blue color (*) represents statistical significances between 15-months muscle transplanted with $FAP^{RFP}$-WT vs. 15-months muscle transplanted with $FAP^{RFP}$-KO.

**Supplementary Figure 4** (r**elated to Main Figure 4)**

(**A**) Flow cytometric isolation strategy to of mononucleated, single, viable/alive (Propidium $iodide^{-}Calcein^{+}$) cells for droplet-based single cell RNA sequencing (scRNA-seq) preparation of total alive cells FACS-isolated from young (2 months) and aged (15 months) TA-muscle of females at 4-dpi.

(**B**) Summary of general features of scRNA-seq experiment of total alive cells isolated from young (2 months) and aged (15 months) TA-muscle of females at 4-dpi.

(**C**) Cell type-scoring of nine subsets derived from unsupervised Louvain clustering (depicted in Main Figure 4B) using marker gene sets (number of genes indicated in each panel; Supplementary Table S3) of cell types associated with regenerating skeletal muscle.

(**D**) Cluster analysis visualization of the top ten most variably expressed genes among the nine identified cell population clusters (listed in Supplementary Table 4S) based on Wilcoxon rank sum testing ($p<0.01$).

(**E**) Enriched terms and pathways of downregulated genes ($p\leq0.05$) by aging in the FAPs subcluster of the total alive cell scRNA-seq dataset. Gene Ontology terms of biological process (BP), cellular component (CC), molecular function (MF), pathway enrichment of reactome database using Metascape gene annotation and analysis tools.

**Supplementary Figure 5** (r**elated to Main Figure 5)**

(**A**) Fluorescence-minus-one (FMO) staining controls and drawn gates used for FACS-isolations and analyses of macrophages in this study with the following surface marker antibody-fluorochrome combinations: CD45-PerCP-Cy5.5; CD11b-PE; F4/80-FITC; Ly6C-PeCy7; CD206-APC.

(**B**) FACS-analysis of total $CD45^{+}CD11b^{+}F4/80^{+}$ macrophages (MCs) normalized to grams of uninjured (uninj) TA-muscle and TA muscle at 4- and 8-dpi comparing young (2 months; white bars) and aged (15 months; blue bars) or *Postn* wildtype (WT; gray bars) and *Postn*$^{-/-}$ (KO; orange bars) male mice (uninj and 8-dpi n=3/group, 4-dpi n=6/group).

(**C**) FACS-analysis of total CD45$^+$CD11b$^+$F4/80$^+$ macrophages (MCs) normalized to grams of uninjured (uninj) TA-muscle and TA muscle at 4- and 8-dpi comparing young (2 months; white bars) and aged (15 months; blue bars) or *Postn* wildtype (WT; gray bars) and *Postn*$^{-/-}$ (KO; green bars) female mice (n=4 in all groups).

(**D**) FACS-analysis of total CD45$^+$CD11b$^+$F4/80$^+$ macrophages (MCs) expressed as percentages of total alive cells for injured TA-muscle at 4- and 8-dpi comparing young (2 months; white bars) and aged (15 months; blue bars) or *Postn* wildtype (WT; gray bars) and *Postn*$^{-/-}$ (KO; orange bars) male mice (4-dpi n=6/group; 8-dpi n=3/group).

(**E**) FACS-analysis of total CD45$^+$CD11b$^+$F4/80$^+$ macrophages (MCs) expressed as percentages of total alive cells for injured TA-muscle at 4- and 8-dpi comparing young (2 months; white bars) and aged (15 months; blue bars) or *Postn* wildtype (WT; gray bars) and *Postn*$^{-/-}$ (KO; green bars) female mice (n=4 in all groups).

(**F**, **G**) Representative density plots and quantification of FACS-analyses of CD45$^+$CD11b$^+$F4/80$^+$Ly6C$^{high}$ and -Ly6C$^{low}$ macrophage frequencies expressed as percentages of total alive cells in injured TA-muscle at 4- and 8-dpi comparing young (2 months; white bars) and aged (15 months; blue bars) or *Postn* wildtype (WT; gray bars) and *Postn*$^{-/-}$ (KO; green bars) female mice (young vs. aged: n=3/group; *Postn*-WT vs. -KO: n=4/group).

(**H**, **I**) Representative density plots and quantification of FACS-analyses of CD45$^+$CD11b$^+$F4/80$^+$CD206$^+$ anti-inflammatory macrophage frequencies expressed as percentages of total alive cells in injured TA-muscle at 4- and 8-dpi comparing young (2 months; white bars) and aged (15 months; blue bars) or *Postn* wildtype (WT; gray bars) and *Postn*$^{-/-}$ (KO; green bars) female mice (young vs. aged: n=3/group; *Postn*-WT vs. -KO: n=4/group).

All animals for experiments were between 2 and 3 months of age, unless indicated otherwise. All data are presented as mean ± standard error of the mean (SEM), statistical significances are *p<0.05, **p<0.01, ***p<0.001.

**Supplementary Figure 6** (r**elated to Main Figure 6)**

**(A)** mRNA expression of pro-inflammatory marker genes *Nos2* (Nitric oxide synthase 2), *Tnfa* (Tumor necrosis factor alpha), *Il1b* (Interleukin 1 beta) and anti-inflammatory marker genes *Cd163* (CD163 antigen) and *Il10* (Interleukin 10) in differentiated bone marrow-derived macrophages (BMDM) exposed to mouse recombinant pro-inflammatory cytokine Interferon-gamma (mrIFNγ; 50 ng/mL; light gray; n=8), mouse recombinant anti-inflammatory cytokine Interleukin-4 (mrIL4; 10 ng/mL; dark gray; n=8), or co-cultured with *Postn*-expressing (FAPs-WT (Postn$^{+/+}$); brown bar; n=11), or *Postn*-deficient (FAPs-KO (Postn$^{-/-}$); blue bar; n=11) FAPs for 48h compared to control BMDM (Ctrl; white bars; n=6) in 3 independent experiments.

**(B)** mRNA expression of genes indicated in panel S6A in differentiated BMDM treated with mouse recombinant Periostin (mrPOSTN) at indicated concentrations (0, 0.25, 0.5, 1.0 µg/mL) for 48h (n=3/group).

**(C)** mRNA expression of genes indicated in panel S6A in differentiated control BMDM (white bar) compared to BMDM 48h-exposed to Cilengitide (10 nM; light gray bar), mrPOSTN (1.0 µg/mL; black bar), mrPOSTN + Cilengitide (dark blue bar), mrIL4 (10 ng/mL; dark gray bar) or rIL4 + Cilengitide (light blue bar) (n=4/group).

**(D)** mRNA expression of pro-inflammatory marker genes *TNFA*, (Tumor necrosis factor alpha), *IL1B* (Interleukin 1 beta), and *CCL2* (C-C Motif chemokine ligand 2) and anti-inflammatory marker genes *CD163* (CD163 antigen), *MRC1* (Mannose receptor c-type 1), and *IL10* (Interleukin 10) in differentiated human THP-1 monocytes treated with human recombinant Periostin (hrPOSTN) at indicated concentrations (0, 0.25, 0.5, 1.0 µg/ml) for 48h (n=3/group).

**(E)** mRNA expression in of genes indicated in panel S6D in differentiated control THP-1 monocytes (white bar) compared to THP-1 monocytes 48h-exposed to Cilengitide (10 nM; light gray), hrPOSTN (1.0 µg/mL; black bar), hrPOSTN + Cilengitide (dark blue bar), hrIL4 (10 ng/mL; dark gray) or hrIL4 + Cilengitide (light blue bar) n=2-3/group; 3 independent experiments).

**(F)** Representative immunofluorescence images (green: CD68, macrophage marker; red: CD206, M2-marker; blue: DAPI, nuclei; scale bar: 20 µm; n=2/group) in untreated, differentiated THP-1 macrophages with solvent control (Vehicle) compared to a 48h treatment with rhPOSTN (1 µg/mL) alone or in combination with Cilengitide (10 nM; rhPOSTN + Ci). Recombinant human IL4 (rhIL4; 10 ng/mL) was used as positive control of M2-polarization.

**(G)** Protein detection and quantification of phosphorylated Ser473-AKT (p-AKT) and total AKT (t-AKT) using Actin as loading control in differentiated THP-1 cells without treatment (white bar) compared to THP-1 cells treated with rhPOSTN (1 µg/mL; gray bars) alone or combined with Cilengitide (10 nM; red bars) for indicated times (n=4/group; 2 independent experiments).

(**H**) Protein detection and quantification of phosphorylated Tyr397-FAK (p-FAK) and total FAK (t-FAK) using Actin as loading control in differentiated THP-1 cells without treatment (white bar) compared to THP-1 cells treated with rhPOSTN (1 µg/mL; gray bars) alone or combined with Cilengitide (10 nM; red bars) for indicated times (n=4; 2 independent experiments).

**Supplementary Figure 7** (r**elated to Main Figure 6)**

(**A**) Mechanical characterization of soft and stiff hydrogels without (control, grey bars) or with mrPOSTN alone (1 µg/mL; red bars), BMDM that were co-treated with mrPOSTN and Cilengitide (10 nM; blue bar), compared to

Cilengitide alone (dark gray bar) for 24h (n=12/group; 2 independent experiments). All data are presented as mean ± standard error of the mean (SEM).

(**B**) Representative immunofluorescence images of F4/80$^{+}$CD86$^{+}$ pro-inflammatory MC-polarization (green: CD86, M1-marker; red: F4/80, macrophage-marker; blue: DAPI, nuclei; scale bar: 206.3 µm) of differentiated BMDM encapsulated in low or high stiffness hydrogels without (Vehicle) compared to Cilengitide alone (10 nM), or mrPOSTN alone (1 µg/mL), or co-treatment with mrPOSTN and Cilengitide (mrPOSTN + Ci) for 24h.

(**C**) Representative immunofluorescence images of F4/80$^{+}$CD206$^{+}$ anti-inflammatory MC-polarization (green: CD206, M2-marker; red: F4/80, macrophage-marker; blue: DAPI, nuclei; scale bar: 206.3 µm) of differentiated BMDM encapsulated in low or high stiffness hydrogels without (Vehicle) compared to Cilengitide alone (10 nM), or mrPOSTN alone (1 µg/ml), or co-treatment with mrPOSTN and Cilengitide (mrPOSTN + Ci), for 24h.

**Supplementary Tables**

**Supplementary Table S1**: Gene lists of microarray analysis of 6-weeks old, young compared to 15-months old, aged FAP FACS-isolated from uninjured hindlimb muscle of male mice. Lists of all detected genes for the individual samples and analyzed data summarizing significantly regulated genes are shown (fold change ≥ -1.5; p≤0.05). Gene annotations and enriched pathway terms were identified using Metascape.

**Supplementary Table S2**: Down-regulated gene list (fold change ≥ -1.5; p value ≤ 0.05) of microfluidics-based scRNA-seq analysis of enriched FAPs isolated form male mice either aged 2 months or 15 months using injured TA muscle at 4-dpi. Among the detected genes, 519 were up- and 1128 gene were downregulated (fold change ≥ 1.5; p value ≤ 0.05). Among the up-regulated genes, computational analysis predicted 70 secreted genes.

**Supplementary Table S3**: Gene list for cell population scoring for supervised clustering of droplet-based scRNA-seq dataset from FACS-sorted viable, non-myofiber cells in regenerating female muscle at 4-dpi (Main Figure 4).

**Supplementary Table S4**: Top 10 scoring genes as visualized in suppl. Figure S4D, organized by the average of fold changes in individual cell transcriptomes from droplet-based scRNA-seq dataset of non-myofiber populations of TA muscle at 4-dpi comparing young and aged female mice. Cell types are based on Louvain clustering (Main Figures 4B and 4C).

**Supplementary Table S5**: Percentages of cell type subsets comparing injured TA of mice either aged 2 months or 15 months using droplet-based scRNA-seq dataset from FACS-sorted viable, non-myofiber cells in regenerating female muscle at 4-dpi (Main Figure 4C).

**Supplementary Table S6**: Differentially expressed genes (DEGs) in defined cell type subsets comparing injured TA (4-dpi) of mice either aged 2 months or 15 months using droplet-based scRNA-seq dataset from FACS-sorted viable, non-myofiber cells in regenerating female muscle at 4-dpi (Main Figure 4).

**Supplementary Table S7**: Summary of Periostin-based cell-cell-interactions (CCI) in published database as reported by Lagger *et al*. in the 'scAgeCom' database.

#### Materials and methods

#### Materials table

| Reagent or resource | Source | Identifier |
|---|---|---|
| **Antibodies** | | |
| Anti-Mouse Ly-6A/E (Sca-1) APC (Clone: D7) | Thermo Fisher Scientific | Cat#17-5981-82;<br>RRID:AB_469487 |
| Anti-Mouse CD45 APC/-eFluor 780 (Clone: 30-F11) | Thermo Fisher Scientific | Cat#47-0451-82;<br>RRID:AB_1548781 |
| Anti-Mouse CD45 FITC (Clone: 30-F11) | Thermo Fisher Scientific | Cat#11-0451-82;<br>RRID:AB_465050) |
| Anti-Mouse CD45 PerCP/Cy5.5 (Clone: 30-F11) | BioLegend | Cat#103131;<br>RRID:AB_893344 |
| Anti-Mouse CD31 (PECAM-1) PE-Cyanine7 (Clone: 390) | Thermo Fisher Scientific | Cat#25-0311-82;<br>RRID:AB_2716949 |
| Anti-Mouse CD31 (PECAM-1) FITC (Clone: 390) | Thermo Fisher Scientific | Cat#11-0311-82;<br>RRID:AB_465012 |
| Anti-Mouse CD11b PE (Clone: M1/70) | Thermo Fisher Scientific | Cat#12-0112-82;<br>RRID:AB_2734869) |
| Anti-Mouse CD11b FITC (Clone: M1/70) | Thermo Fisher Scientific | Cat#11-0112-82;<br>RRID:AB_464935 |
| Anti-Mouse F4/80 FITC (Clone: BM8) | Thermo Fisher Scientific | Cat#11-4801-82;<br>RRID:AB_2637191 |
| Anti-Mouse Integrin α7 PE (Clone: 3C12) | Miltenyi Biotec | Cat#130-120-812;<br>RRID:AB_2784388 |
| Anti-Mouse Ly-6C PE-Cyanine7 (Clone: HK1.4) | BioLegend | Cat#128017;<br>RRID:AB_1732093 |
| Anti-Mouse CD206 (MMR) APC (Clone: C068C2) | BioLegend | Cat#141707;<br>RRID:AB_10896057 |

| | | |
|---|---|---|
| TruStain FcX™ (anti-mouse CD16/32) (Clone: 93) | BioLegend | Cat#101319;<br><br>RRID:AB_1574973 |
| Rat anti-F4/80 | Abcam | Cat#ab6640;<br><br>RRID:AB_1140040) |
| Rabbit Anti-CD206 (MMR) | Abcam | Cat#ab64693;<br><br>RRID:AB_1523910 |
| Rabbit Anti-CD86 | Abcam | Cat#ab239075,<br><br>RRID:AB_2927417 |
| Mouse Anti-CD68 | Abcam | Cat#ab955;<br><br>RRID:AB_307338 |
| Rat Anti-Laminin B2, clone A5 | Millipore | Cat#05-206;<br><br>RRID:AB_309655 |
| Mouse anti-eMHC | DHSB | Cat#F1.652;<br><br>RRID:AB_528358 |
| Rabbit anti-POSTN | Novus Biologicals | Cat#NBP1-30042;<br><br>RRID:AB_1968578 |
| Rabbit anti-DESMIN | Abcam | Cat#ab32362;<br><br>RRID:AB_731901 |
| Rabbit anti-Ki67 | Abcam | Cat#15580;<br><br>RRID:AB_443209 |
| Goat anti-PDGFRα | R&D systems | Cat#AF1062;<br><br>RRID:AB_2236897) |
| Rabbit anti-RFP (tdTomato) | Abcam | Cat#ab62341;<br><br>RRID:AB_945213 |
| Rabbit anti-AKT | Cell Signaling Technology | Cat#9272;<br><br>RRID:AB_329827 |
| Rabbit anti-phospho AKT (Ser473) | Cell Signaling Technology | Cat#4060;<br><br>RRID:AB_2315049 |

| | | |
|---|---|---|
| Rabbit anti-FAK | Cell Signaling Technology | Cat#3285;<br>RRID:AB_2269034 |
| Rabbit anti-phospho FAK (Tyr397) | Cell Signaling Technology | Cat#3283;<br>RRID:AB_2173659 |
| Mouse anti-ACTIN | Cell Signaling Technology | Cat#41185;<br>RRID:AB_3065071 |
| Mouse anti-eMHC | DHSB | Cat#F1.652;<br>RRID:AB_528358 |
| Alexa Fluor 488 goat anti-rabbit | Thermo Fisher Scientific | Cat#A-11008;<br>RRID:AB_143165 |
| Alexa Fluor 488 goat anti-mouse | Abcam | Cat#ab150113;<br>RRID:AB_2576208 |
| Alexa Fluor 488 chicken anti-goat | Thermo Fisher Scientific | Cat#A-21467;<br>RRID:AB_2535870 |
| Alexa Fluor 555 goat anti-rat | Thermo Fisher Scientific | Cat#A-21434;<br>RRID:AB_2535855 |
| Alexa Fluor 568 goat anti-rabbit | Abcam | Cat#ab175471;<br>RRID:AB_2576207) |
| Alexa Fluor 594 donkey anti-rabbit | Thermo Fisher Scientific | Cat#A-21207;<br>RRID:AB_141637 |
| IRDye 680RD Goat anti-Mouse IgG – for western blot | LI-COR Biosciences | Cat#925-68070;<br>RRID:AB_2651128 |
| IRDye 800CW Donkey anti-Rabbit IgG – for western blot | LI-COR Biosciences | Cat#925-32213,<br>RRID:AB_2715510 |
| **Chemicals, Peptides, and Recombinant Proteins** | | |
| Glycerol | Sigma-Aldrich | Cat#G2025 |
| Collagenase A | Sigma-Aldrich | Cat#10103586001 |
| Dispase II | Thermo Fisher Scientific | Cat#17105041 |

| | | |
|---|---|---|
| Calcein Blue AM Viability Dye | eBioscience | Cat#65-0855-39 |
| Propidium Iodide (PI) | Sigma-Aldrich | Cat#P4170 |
| Roti-Histofix 4 % | Carl Roth | Cat#P087.3 |
| Tragacanth | Sigma-Aldrich | Cat#G1128 |
| Triton™ X-100 | Sigma-Aldrich | Cat#X100 |
| Fluoromount-G | eBioscience | Cat#00-4958-02 |
| 4′,6-diamidino-2-phenylindole (DAPI) | BioLegend | Cat#422801 |
| X-β-Gal | Carl Roth | Cat#2315.4 |
| Matrigel-Matrix | Corning | Cat#356231 |
| DMEM/high glucose | Pan Biotech | Cat#P04-03590 |
| Ham´s F10 Medium | Pan Biotech | Cat#P04-12500 |
| Penicillin /Streptomycin | Pan Biotech | Cat#P06-07100 |
| Fetal Bovine Serum | Biochrom | Cat#S0115 |
| Basic fibroblast growth factor | Sigma-Aldrich | Cat#F0291 |
| DMEM/low glucose | Pan Biotech | Cat#P04-01550 |
| Horse Serum, New Zealand origin | Thermo Fisher Scientific | Cat#16050122 |
| MCDB201 Media | Sigma-Aldrich | Cat#M6770 |
| Insulin-transferrin-selenium (ITS) mix | Sigma-Aldrich | Cat#I3146 |
| Linoleic acid-Albumin | Sigma-Aldrich | Cat#L9530 |
| Dexamethasone | Sigma-Aldrich | Cat#D-4902 |
| L-Ascorbic acid 2-phosphate | Sigma-Aldrich | Cat#A8960 |
| BODIPY™ 493/503 | Thermo Fisher Scientific | Cat#D3922 |
| DMEM/F12 (1:1) | Pan Biotech | Cat#P04-41250 |
| Recombinant Murine Macrophage Colony Stimulating Factor | PeproTech | Cat#315-02; GenPept: P07141 |
| Recombinant Mouse Periostin | R&D systems | Cat#2955-F2-050; GenPept: NP_056599 |

| | | |
|---|---|---|
| Recombinant Human Periostin | R&D systems | Cat#3548-F2-050; GenPept: Q15063 |
| Recombinant Murine IL-4 | PeproTech | Cat#214-14; GenPept: P07750 |
| Recombinant Human IL-4 | PeproTech | Cat#200-04; GenPept: P05112 |
| Recombinant Murine IFNγ | PeproTech | Cat#315-05; GenPept: P051580 |
| Cilengitide trifluoroacetate | Selleckchem | Cat#S7077; CAS: 199807-35-7 |
| RPMI-1640 | Pan Biotech | Cat#P04-18047 |
| β-Mercaptoethanol | Sigma-Aldrich | Cat# M6250 |
| Phorbol 12-myristate 13-acetate | Sigma-Aldrich | Cat#P8139 |
| Protease Inhibitor Cocktail | Sigma-Aldrich | Cat#P8340 |
| Phosphatase Inhibitor Cocktail 2 | Sigma-Aldrich | Cat#P5726 |
| Phosphatase Inhibitor Cocktail 3 | Sigma-Aldrich | Cat#P0044 |
| Bovine Serum Albumin (BSA) Fraction V | Pan Biotech | Cat#P06-1391500 |
| Normal goat serum | Abcam | Cat#ab7481 |
| Intercept® (TBS) Blocking Buffer | LI-COR Biosciences | Cat#927-60003 |
| TRIzol® Reagent | Fisher Scientific | Cat#12034977 |
| 1 µm Transwell inserts | Corning | Cat#353104 |
| SeqAmp™ DNA Polymerase | Takara Bio, Inc. | Cat#: 638504 |
| Protector RNase Inhibitor | Sigma Aldrich | Cat#: 3335399001 |
| 1X TE Buffer | Thermo Fisher Scientific | Cat#: PN12090015 |
| SPRIselect | Beckman Coulter Life Sciences | Cat#: B23318 |
| Agencourt AMPure XP Beads | Beckman Coulter Life Sciences | Cat#: A63880 |

| **Software and algorithms used** | | |
|---|---|---|
| Cellpose | cellpose.org | RRID:SCR_021716 |
| Metascape | metascape.org | RRID:SCR_016620 |
| ImageJ | imagej.net | RRID:SCR_003070 |
| Fiji | fiji.sc | RRID:SCR_002285 |
| CFX Manager | Bio-Rad | RRID:SCR_017251 |
| Odyssey CLx | LI-COR | RRID:SCR_014579 |
| FlowJo™ v10.5.3 | BD Biosciences | RRID:SCR_008520 |
| Matlab | mathworks.com | RRID:SCR_001622 |
| STAR (version 2.6.0a) | tabit.ucsd.edu/sdec | RRID:SCR_005622 |
| featureCounts (version 1.6.2) | bioinf.wehi.edu.au/featureCounts | RRID:SCR_012919 |
| Seurat (version 3) | satijalab.org/seurat/get_started.html | RRID:SCR_016341 |
| Cell Ranger (V4.0.0) | 10xgenomics.com | RRID:SCR_017344 |
| GraphPad Prism (version 9.4.1) | graphpad.com | RRID:SCR_002798 |
| Cell lines used | | |
| THP-1 | ATCC | Cat#TIB-202; RRID:CVCL_0006 |
| **Critical Commercial Assays** | | |
| Periostin ELISA | Biomedica | Cat#BI-20433 |
| Pierce BCA Protein Assay Kit | Thermo Fisher Scientific | Cat#23225 |
| RNAqueous Micro Kit | Thermo Fisher Scientific | Cat#AM1931 |
| Affymetrix Mouse Exon 1.0 ST Array | Thermo Fisher Scientific | Cat#901168 |
| RNA Miniprep Kit | Zymo Research | Cat#1065 |
| High Capacity cDNA Reverse Transcription Kit | Fisher Scientific | Cat#10186954 |
| Maxima SYBR Green/ROX qPCR Master Mix | Fisher Scientific | Cat#11893913 |
| Agilent High Sensitivity DNA Kit | Agilent Technologies | Cat#: 5067- 4626 |

| | | |
|---|---|---|
| Agilent DNA 1000 Kit | Agilent Technologies | Cat#: 5067-1504 |
| SMART-Seq® v4 Ultra® Low Input RNA Kit for the Fluidigm® C1™ System | Takara Bio, Inc. | Cat#: 635025 |
| C1™ Single-Cell mRNA Seq HT 10-17 µm IFC | Fluidigm | Cat#: PN101-4982 |
| Kit, C1™ Single-Cell Reagent Kit for mRNA Seq | Fluidigm | Cat#: PN100-6201 |
| Quant-iT™ PicoGreen™ dsDNA Assay Kit | Invitrogen | Cat#: P7589 |
| Nextera XT DNA Library Preparation Kit | Illumina | Cat#: FC-131-1096 |
| Nextera XT Indexing Kit v2 Set A | Illumina | Cat#: FC-131-2001 |
| Nextera XT Indexing Kit v2 Set B | Illumina | Cat#: FC-131-2002 |
| Chromium Next GEM Single Cell 3' Kit v3.1, 4 rxns | 10X Genomics | Cat#: PN1000269 |
| Chromium Next GEM Chip G Single Cell Kit | 10X Genomics | Cat#: PN-1000127 |
| Agilent High Sensitivity DNA Kit | Agilent Technologies | Cat#: 5067- 4626 |
| **Deposited Data** | | |
| scRNA-sequencing; microfluidics-based dataset of FAPs (https://0-www-ncbi-nlm-nih-gov.brum.beds.ac.uk/geo/query/acc.cgi?acc=GSE247415) | NCBI GEO-Database (Gene Expression Omnibus) | Accession number: GSE247415 |
| scRNA-sequencing; droplet-based dataset of all alive, non-myofiber cells (https://0-www-ncbi-nlm-nih-gov.brum.beds.ac.uk/geo/query/acc.cgi?acc=GSE247313) | NCBI GEO-Database (Gene Expression Omnibus) | Accession number: GSE247313 |
| **Experimental Models: Organisms/Strains** | | |
| Mouse, strain: C57BL/6J | The Jackson Laboratory | RRID:IMSR_JAX:000664 |
| Mouse, strain: B6N(Cg)-Postntm1.1(KOMP)Vlcg/J | The Jackson Laboratory | RRID:IMSR_JAX:024186 |
| Mouse, strain: B6.129(Cg)-Gt(ROSA)26Sortm4(ACTB-tdTomato,-EGFP)Luo/J | The Jackson Laboratory | RRID:IMSR_JAX:007576 |
| **Oligonucleotides** | | |

| | | |
|---|---|---|
| Fwd: CGGGAAGAACGAATCATTACA<br>Rev: ACCTTGGAGACCTCTTTTTGC | This manuscript | Postn: mouse, NM_015784.3 |
| Fwd: CGACGAGGAAGGAGACAAGA<br>Rev: CGGGTTCTGATTCCACATCT | This manuscript | Pax7: mouse, NM_011039 |
| Fwd: CAGTGAATGCAACTCCCACA<br>Rev: GAGCAAATGATCTCCTGGGT | This manuscript | Myog: mouse, NM_031189 |
| Fwd: GCTACCCAAGGTGGAGATCCT<br>Rev: GGCGGTGTCGTAGCCATT | This manuscript | Myod: mouse, NM_010866 |
| Fwd: CTGTGTGCAATGCCTATGAGA<br>Rev: CTGGAGGGTATTGAAGAGCCG | This manuscript | Plin1: mouse, NM_001113471 |
| Fwd: GATGCCTTTGTGGGAACCT<br>Rev: CTGTCGTCTGCGGTGATTT | This manuscript | Fabp4: mouse, NM_024406 |
| Fwd: GTGCTCCTGGTATTGCTGGT<br>Rev: GGCTCCTCGTTTTCCTTCTT | This manuscript | Col1a1: mouse, NM_007742 |
| Fwd: ACGTAAGCACTGGTGGACAGA<br>Rev: GAGGGCCATAGCTGAACTGA | This manuscript | Col3a1: mouse, NM_009930.2 |
| Fwd: CCACAGCATTGAGGAGTTTG<br>Rev: ACAGCTCATCATTTGGCTCA | This manuscript | Mrc1: mouse, NM_008625.2 |
| Fwd: GTTCTCAGCCCAACAATACAAGA<br>Rev: GTGGACGGGTCGATGTCAC | This manuscript | Nos2: mouse, NM_ 010927.3 |
| Fwd: GTTCTCAGCCCAACAATACAAGA<br>Rev: GTGGACGGGTCGATGTCAC | This manuscript | Tnfa: mouse, NM_ 013693 |
| Fwd: AGTTGACGGACCCCAAAAG<br>Rev: AGCTGGATGCTCTCATCAGG | This manuscript | Il1b: mouse, NM_ 008361.3 |
| Fwd: TCCACACGTCCAGAACAGTC<br>Rev: CCTTGGAAACAGAGACAGGC | This manuscript | Cd163: mouse, NM_001170395.1 |
| Fwd: CAGAGCCACATGCTCCTAGA<br>Rev: TGTCCAGCTGGTCCTTTGTT | This manuscript | Il10: mouse, NM_ 010548.2 |

| | | |
|---|---|---|
| Fwd: TTTGGGCATCACCACGAAAA<br>Rev: GGACACCCTCCAGAAAGCGA | This manuscript | Arbp: mouse,<br>NM_ 007475.5 |
| Fwd: TGGAGCTGGCCGAGGAG<br>Rev: AGCAGGCAGAAGAGCGTGG | This manuscript | TNFA: human,<br>NM_ 000594.4 |
| Fwd: GTGGCAATGAGGATGACTTGTTCT<br>Rev: TGTAGTGGTGGTCGGAGATTCG | This manuscript | IL1B: human,<br>NM_ 000576.3 |
| Fwd: AAACTGAAGCTCGCACTCTCGC<br>Rev: AGGTGACTGGGGCATTGATTG | This manuscript | CCL2: human,<br>NM_ 002982.4 |
| Fwd: AGCAGACTACTCCAACATCC<br>Rev: TGGCACAGTTGTCTCTATCC | This manuscript | CD163: human,<br>NM_ 004244.6 |
| Fwd: GGGTTGCTATCACTCTCTATGC<br>Rev: TTTCTTGTCTGTTGCCGTAGTT | This manuscript | MRC1: human,<br>NM_ 002438.4 |
| Fwd: GACTTTAAGGGTTACCTGGGTTG<br>Rev: TCACATGCGCCTTGATGTCTG | This manuscript | IL10: human,<br>NM_001101.5 |
| Fwd: ATTGCCGACAGGATGCAGAA<br>Rev: GCTGATCCACATCTGCTGGAA | This manuscript | ACTB: human,<br>NM_000572.3 |

#### Methods

##### Mouse models

All procedures were approved by the local ethics committee for animal welfare of the State Office of Environment, Health, and Consumer Protection (Federal State of Brandenburg, Germany). Animals were housed in a controlled environment (20±2 °C, 12 h/12 h light/dark cycle) and maintained on a standard diet (Ssniff, Soest, Germany). Typically, male and female mice were used in the experiments separately at the indicated ages and time points and are presented in separate figure items. The following mouse strains were obtained from The Jackson Laboratory (Bar Harbor, ME, USA) and maintained as local colonies: C57BL/6J, B6N(Cg)-Postntm1.1(KOMP)Vlcg/J (*Postn* mutant allele), B6.129(Cg)-Gt(ROSA)26Sortm4(ACTB-tdTomato,-EGFP)Luo/J (mTmG-reporter) strain. For some experiments, the B6N(Cg)-Postntm1.1(KOMP)Vlcg/J mouse strain was intercrossed with the mTmG-reporter mouse strain that constitutively expresses the membrane-bound, red fluorescent protein tdTomato to generate B6N(Cg)-Postntm1.1(KOMP)Vlcg/J wildtype and knockout allele carriers expressing the tdTomato red fluorescent protein constitutively in transplanted fibro adipogenic progenitor cells (WT-FAP$^{RFP}$ and KO-FAP$^{RFP}$) used for transplantation experiments.

**Muscle injury and FAP transplantation**

Muscle injury was induced by intramuscular injections of 25 µL of 50% glycerol (v/v) (Sigma-Aldrich) in PBS divided in two injections of 12.5 µL each into the tibialis anterior (TA) muscle in mice anesthetized with isoflurane with an analgesic injection of 0.1 mg/kg of buprenorphine 30 min prior to injection. At the indicated days post injection (dpi), mice were killed and TA-muscles were collected for subsequent analyses. For intramuscular FAP transplantations, 20,000 FACS-purified FAPs pooled from two mice were freshly isolated from uninjured hindlimb muscles. Cells were resuspended in PBS, counted, adjusted to a volume 25 µL and directly injected in injured TA-muscle one day after the glycerol injection in two injections of 12.5 µL each into the injury area. After 8- and 14-dpi, FAPs-transplanted TA-muscles were collected for subsequent analyses. Injections were performed under sterile conditions using Hamilton syringes (CS-Chromatography Service GmbH) for FAP transplants and 29-G insulin syringes (BD Biosciences) for glycerol.

**Flow cytometry and fluorescence-activated cell sorting (FACS)**

Flow cytometry and cell sorting were performed on a FACS Aria III cell sorter (BD Biosciences) and analyzed using FlowJo™ Software (Tree Star). Hindlimb muscles (gastrocnemius, quadriceps, extensor digitorum longus, soleus and tibialis anterior muscles) or injured tibialis anterior muscles were isolated, minced and digested in high-glucose DMEM medium containing 2.5 mg/mL of Collagenase A (Sigma-Aldrich) for 45 min at 37 Cº with shaking. 2 U/mL of Dispase II (Thermo Fisher Scientific) were added for 30 minutes to the muscle lysates. Muscles slurries were passed 10 times through a 20 G syringe and a 70 µm cell strainer and centrifugated at 1200 rpm for 5 minutes at 4 ºC. The pellet was re-suspended in Ammonium Chloride Potassium (ACK) lysis buffer/ to eliminate red blood cells and centrifuged again at 1200 rpm for 5 minutes at 4 ºC. Cells were re-suspended in sorting buffer consisting of 100 µl Hank´s balanced salt solution (HBSS) containing 0.4% FBS (Merck Biochrom; sorting buffer) and blocked using anti-mouse CD16/32 FcγR (BioLegend) for 10 minutes, washed and re-suspended in 100 µl of fluorophore-conjugated antibodies in sorting buffer for 30 minutes at 4 ºC. The applied FACS antibodies can be found in the materials table. Living cells were gated for the accumulation of calcein blue (1:1,000 dilution; stock of 1 mg in 215 mL DMSO) and the exclusion of propidium iodide (PI; 1:1,000 dilution; stock of 1 µg/mL in distilled water) fluorescence.

**Primary cell culture (FAPs, MuSCs and BMDM)**

For in vitro experiments, FAPs ($CD45^{-}CD11b^{-}CD31^{-}SCA1^{+}$) and MuSCs ($CD45^{-}CD11b^{-}CD31^{-}SCA1^{-}ITGA7^{+}$) primary cells were sorted by FACS (described above) and plated on 24- or 48-well plates, respectively, coated with 2% Matrigel (Corning) for culture expansion except where stated otherwise. For FAP cultivation, a combined culture medium of 60% DMEM-low glucose (Pan Biotech) and 40 % MCDB201 (Sigma-Aldrich) was supplemented with 1% Penicilin/S Streptomycin (P/S; Pan Biotech). 2 % Fetal bovine serum (FBS) (Merck Biochrom), 1x insulin-transferrin-selenium (ITS) mix, 1x linoleic acid conjugated to bovine serum albumin (BSA), 1 nM dexamethasone, 0.1 mM L-ascorbic acid 2-phosphate and 5 ng/mL recombinant basic fibroblast growth factor (bFGF) (all from Sigma-Aldrich) were added and used as FAP growth medium for expansion during 5 days with medium replacements every 2 days. For spontaneous adipogenesis, after expansion, FAPs were left to spontaneously differentiate for 6 days in FAP-growth medium with medium replacement at day 3, but without the addition of bFGF. For adipocyte staining, cells were fixed in 4 % Formaldehyde (Carl Roth) for 10 minutes, then incubated with BODIPY 493/503 (Thermo Fisher Scientific) in a 1:2,500 dilution in Phosphate-buffered saline (PBS; BODIPY prepared from 2 µM stock in DMSO) and counterstained with 300 nM of 4′,6-diamidino-2-phenylindole (DAPI; BioLegend) from 5 minutes in PBS.

Following FACS-isolation, MuSCs were resuspended in myogenic growth medium containing DMEM-high glucose and Ham’s F10 (50 % v/v; Pan Biotech), 20 % FBS (Merck Biochrom), 1 % P/S (Pan Biotech) and 5 ng/mL bFGF (Sigma-Aldrich). After 5 days of expansion with medium replacement every 2 days, MuSCs were differentiated into myofibers by switching to myogenic differentiation medium containing DMEM-low glucose (Pan Biotech), 2 % horse serum (Thermo Fisher Scientific) and 1 % P/S (Pan Biotech) medium for 5 days with medium replacement every other day. Differentiated myofibers were fixed in 4 % Formaldehyde (Carl Roth) for 10 minutes and washed 3 times for 5 minutes with PBS at room temperature (RT). Permeabilization solution with 0.1 % Triton X-100 (Sigma Aldrich) in PBS was applied for 10 minutes and washed for 5 minutes with PBS before blocking solution with 3 % BSA (Pan Biotech) in PBS was added for 1 hour at RT. After blocking, differentiated fibers were stained using a rabbit polyclonal anti-Desmin (Abcam) diluted 1:100 in blocking solution, overnight at 4 °C. On the following day, after washing, cells were incubated for 1 hour with the corresponding secondary antibody (see Materials Table) diluted 1:500 in blocking solution and nuclei were counter-stained with 300 nM DAPI in PBS for 5 minutes. Fusion index was calculated as the percentage of nuclei in desmin positive fibers per total number of fibers. Cells and stainings were assessed in a BZ900 Fluorescence Microscope (Keyence).

For bone marrow derived macrophage (BMDM) assays, bone marrow was flushed from the femurs and tibiae of 8 to 12 weeks-old C57BL/6J mice using a 26 G needle and ice-cold, sterile PBS. Marrow was dispersed by aspirating it through a 19 G needle twice and centrifugated for 5 minutes at 4ºC. Once re-suspended in macrophage differentiation medium containing DMEM/F12 (50 % v/v) (Pan Biotech), 10 % FBS (Merck Biochrom), 1 % P/S (Pan Biotech) and supplemented with 50 ng/mL of recombinant murine macrophage colony stimulating factor (m-CSF) (PeproTech), BMDM were incubated for 7 days in a 150 mm plate with medium replacements six hours after seeding, to initially eliminate non-attached red blood cells, and debris and on day 3. All primary culture experiments were performed at 37 ºC and 5 % $CO_2$.

**Co-culture assays**

For direct-contact co-cultures of FAPs and MuSCs, a total of 15,000 cells was seeded in a 1:1 ratio, 7,500 each, after 3 days of expansion in 48-well plates coated with 2% Matrigel (Corning) and cultured in myogenic differentiation medium for 5 days. Next, cells were fixed (4% formaldehyde, 10 minutes), washed, blocked (3 % BSA in PBS, 1 hour at RT) and co-stained overnight at 4ºC using a goat polyclonal anti-PDGRα antibody (R&D Systems) and a rabbit polyclonal anti-Desmin (Abcam), both diluted 1:100. After washing, cells were incubated for 1 hour with the corresponding secondary antibody diluted 1:500 in blocking solution and nuclei were counter-stained with 300 nM DAPI in PBS. Stainings were visualized in a BZ900 Fluorescence Microscope (Keyence). Images were obtained from 3 independent wells per condition. ImageJ software[1] was used to manually determine fiber lengths of individual fibers along with fusion index determined as the percentage of fibers with >3 $DAPI^+$ nuclei and normalized to total number of fibers.

For trans-well co-culture assays, BMDM were differentiated for 7 days (described above), and $2x10^5$ cells were seeded in 24-well plates in macrophage differentiation medium. Next, 50,000 FAPs isolated from the hindlimb muscles and expanded for 5 days were seeded into 24-well inserts with 1.0 µm mesh (Corning) in FAPs growth medium for 4-6 hours to allow cells to attach. Inserts were then washed gently and macrophage differentiation medium was added. Inserts were added to BMDM-containing wells and co-cultured with BMDM for 48 hours before harvesting of BMDM-wells for gene expression analysis. 50 ng/mL of mouse recombinant interferon gamma (mrIFNγ) or 10 ng/mL of recombinant murine interleukin 4 (mrIl4; both from PeproTech) were used to induce pro- and anti-inflammatory macrophage polarization, respectively, as control conditions.

**Proliferation assays in primary FAPs and MuSCs**

To asses MuSC and FAP proliferation, 5,000 cells were cultured in 96-well plates coated with 2% Matrigel (Corning) in myogenic or FAP growth medium, respectively, alone of with addition of 0.5 µg/mL of mouse recombinant Periostin (mrPOSTN; R&D systems) added for 48 hours. After incubations, cells were fixed in 4% formaldehyde for 10 minutes and washed 3 times for 5 minutes with PBS at RT. Permeabilization solution (0.1% Triton X-100 in PBS) was applied for 10 minutes and washed for 5 minutes with PBS before block solution using 3 % BSA in PBS was added for 1 hour at RT. After blocking, cells were stained overnight at 4ºC using rabbit polyclonal anti-Ki67 antibody (Abcam) diluted at 1:100 in blocking solution. After washing, cells were incubated 1 hour at RT with the corresponding secondary antibody (see Materials Table) diluted 1:500. Nuclei were counter-stained with 300 nM DAPI in PBS for 5 minutes. Staining was visualized in a BZ900 Fluorescence Microscope (Keyence) and random images of Ki67 positive cells were quantified at least from 3 independent wells per condition and normalized to total number of cells.

**THP-1 cell culture**

The human-derived monocytic THP-1 cell line (ATCC) was cultured in RPMI-1640 (Pan Biotech), 10 % FBS (Biochrom), 1 % P/S (Pan Biotech) and supplemented with 0.05 mM β-Mercaptoethanol (Sigma-Aldrich) in 75 $cm^2$ culture T-flasks at 37 ºC and 5 % $CO_2$. Cells were grown to a density of 2-8 x $10^5$ and used for experiments between passages 8 to 12. For differentiation to a macrophage-like phenotype, THP-1 were seeded in 24- or 48-well plates and incubated with 200 nM of phorbol 12-myristate 13-acetate (PMA; Sigma-Aldrich) for 48 hours. Following differentiation, PMA-containing medium was replaced with fresh growth medium for 24 hours before treatments.

**BMDM and THP-1 treatments**

After differentiation of BMDM and THP-1 to macrophages as described above, $5x10^5$ BMDM and THP-1 cells were seeded into 24-well plates in their corresponding medium with the addition of 0.25, 0.5 or 1 µg/mL mouse or human recombinant Periostin (mrPOSTN, hrPOSTN) (R&D systems) for 48 hours. For inhibition of Periostin-dependent signaling via integrins, 1 µg/mL of mrPOSTN (BMDM) or hrPOSTN (THP-1, both from R&D Systems) were used either alone or in combination with 10 nM Cilengitide (Selleck Chemicals GmbH) for 48 hours. 10 ng/mL of recombinant murine or human interleukin 4 (mrIl4 or hrIL4; both from PeproTech) were used as controls to induce an anti-inflammatory (M2-like) phenotype BMDM and THP-1 macrophages, respectively. After 48 hours of incubation, both cell types were used for gene expression or immunofluorescence analyses. For signaling analyses, differentiated

BMDM and THP-1 cells were overnight-incubated in serum-free medium. Cells were then washed with PBS and incubated with 1 µg/mL of mrPOSTN (BMDM) or hrPOSTN (THP-1; both from R&D Systems) alone or combined with 10 nM Cilengitide (Selleck Chemicals GmbH) in serum-free medium for 15, 30 and 60 minutes before harvesting.

**Quantitative real-time PCR**

RNA was extracted from frozen muscles or freshly harvested cells using TRIzol reagent (Fisher Scientific) and purified using commercially available RNA Miniprep Kits (Zymo Research). RNA was transcribed into cDNA with a high capacity cDNA reverse transcription kit (Fisher Scientific). Quantitative real-time PCR (qPCR) was performed using the Biorad CFX384 Real-Time System (Biorad) with Maxima™ SYBR™ Green/ROX 2x qPCR Master Mix (Thermo Fisher Scientific). Primers were designed as intron spanning sequences to specifically amplify cDNA while excluding potential contaminating genomic DNA. mRNA expression was calculated relative to the mRNA expression of reference gene *Arbp* in mouse and *ACTB* in human samples, respectively.

**Protein detection by Western blotting**

For protein analysis, snap-frozen muscle tissue samples and freshly harvested macrophage (BMDM and THP-1) cell cultures were homogenized in RIPA buffer supplemented with protease and phosphatase inhibitor cocktails (Sigma-Aldrich), each diluted 1:100. FAP cell culture supernatants were directly diluted in RIPA buffer (1:5). Protein concentrations were determined with a Pierce BCA protein assay kit (Thermo Fisher Scientific) and equal amounts of protein were diluted in Laemmli sample buffer supplemented with β-mercaptoethanol (1:1000), then boiled at 37 °C for 5 minutes. Protein separation was performed using 10 % SDS-PAGE and subsequently transferred onto PVDF membranes. After blocking using the commercially available Intercept-TBS blocking buffer (LI-COR Biosciences), membranes were incubated with a rabbit polyclonal anti-Periostin (Novus Biologicals), rabbit polyclonal anti-AKT, rabbit polyclonal anti-phospho AKT (Ser473), rabbit polyclonal anti-FAK, rabbit polyclonal anti-phospho FAK (Tyr397) (all diluted 1:1,000 in blocking buffer; overnight at 4Cº) for specific protein detections, and mouse monoclonal anti-Actin (diluted 1:5,000 in blocking buffer; 1 hour at RT) as loading control (all from Cell Signaling Technology). Membranes were then washed and incubated with their corresponding secondary antibodies for 1 hour at RT (see Materials Table). Detection was performed using the Odyssey CLx imaging system (LI-COR Biosciences), signals were quantified using ImageJ software[1], and normalized to Actin loading control.

**Histology, immunofluorescence and image analysis of muscle sections**

For paraffin embedding, muscles were dissected and fixed overnight in 4% formaldehyde (Carl Roth) at 4 ºC, subsequently dehydrated, and embedded in paraffin sections before 2 µm slices were prepared. For TA muscle, uninjured or injury areas were sectioned in the middle region of the muscle at its maximum diameter and consecutive sections were collected every 250 µm for hematoxylin and eosin (H&E) or sirius red staining according to standard histological protocols. Adipocyte infiltration areas within H&E staining of injured muscle were quantified using ImageJ software[1] and normalized to total regeneration area. Collagenous matrix deposition was stained with sirius red and fibrosis areas were quantified using ImageJ software[1] and available plugins "MRI fibrosis tool" and "Color Deconvolution" and normalized to total regeneration area in injured TA or total tissue section area in uninjured muscles. For cryo-sections, TA muscle was collected and embedded in 6 % (w/v) gum tragacanth (Sigma-Aldrich) dissolved in distilled water, and snap frozen in ice-cold isopentane pre-cooled to −160°C. Muscle sections were prepared at 10 µm thickness from regenerating areas and collected onto Superfrost Plus adhesion slides (Thermo Fisher Scientific) and stored at −80°C for immunofluorescence analyses. For laminin (Millipore), eMHC (DHSB) and $RFP^{tdTom}$ (Abcam) immunofluorescence, cryo-sections were thawed for 10 minutes and blocked for 1 hour at RT in blocking solution containing 3 % BSA (w/v, Pan Biotech) and 5 % normal goat serum (v/v; Abcam) in PBS. Slides were stained overnight at 4ºC in a humidified chamber using monoclonal anti-laminin B5 antibody produced in rat (Millipore), the monoclonal anti-eMHC produced in mouse (DSHB) and the polyclonal anti-RFP produced in rabbit diluted at 1:200, 1:80 and 1:100 in blocking solution, respectively. For macrophage and ECM-component immunofluorescence, cryo-sections were allowed to thaw-dry for 10 minutes, sections were subsequently fixed with 1 % formaldehyde, washed briefly and permeabilized with 0.3 % Triton X-100 (Sigma-Aldrich) in PBS for 10 minutes. Afterwards, slices were blocked for 1 hour at RT in blocking solution. Rat monoclonal anti-F4/80 (Abcam), rabbit polyclonal anti-CD206 (Abcam) and rabbit polyclonal anti-POSTN (Novus Biologicals) antibodies diluted at 1:100 in blocking solution were incubated overnight at 4ºC in a humidified chamber. After primary antibody incubation and washing, sections were incubated for 1 hour at RT with the corresponding secondary antibody (see Materials Table) diluted 1:500 and nuclei were counterstained with 300 nM DAPI (BioLegend) in PBS for 5 minutes. After washing, sections were mounted with Fluoromount G (eBioscience) and stainings were visualized using a BZ900 Fluorescence Microscope (Keyence) or confocal microscope (TCS SP8 X, Leica Microsystems) for higher magnification. Collagen and laminin were red-stained to delimit muscle fibers from the extracellular matrix. The CellPose deep-learning algorithm was used to automatically segment individual fibers[2]. Cross sectional area fiber size from 3 different midbelly (uninjured) or in post-injury areas of TA sections per mouse were analyzed with FIJI/ImageJ, plugin LabelsToROIs, as described before[3]. The number of positive macrophages co-localized with CD206 as M2 macrophage marker and eMHC positive fibers were determined manually by counting of positively stained cells in injury areas of muscle sections using the ImageJ software[1]. For X-Gal staining, whole-mount gastrocnemius and soleus muscles were fixed in 4% formaldehyde for 1h at 4Cº, rinsed four times for 30 minutes in rinse buffer (PBS, 2 mM $MgCl_2$, 0.02 % Nonidet P-40, 0.01 % Na-Deoxycholate; all from Sigma-Aldrich) and were incubated overnight with X-gal (2 mM $MgCl_2$, 0.02 % Nonidet P-40, 0.01 % Na-Deoxycholate, 5 mM Potassium ferrocyanide, 1 mg/mL X-β-gal; all form Carl Roth) to enable the β-galactosidase reporter reaction at 37°C in the dark. Subsequently, tissues were washed in PBS, photographed, dehydrated, and embedded in paraffin. 2 µm slices were collected and counterstained with H&E for microscopic analysis of staining distribution.

**Transcriptomic analyses of microarray and single cell RNA-sequencing (scRNA-seq) data**

Microarray expression analysis was conducted on FACS-enriched FAPs ($CD45^-CD11b^-CD31^-SCA1^+$) derived from pooled hind limb skeletal muscles of young (6 weeks) and old (15 months) C57BL/6J male mice without injury using an analysis strategy described before[4]. Primary FAPs were purified by FACS and cell pellets were immediately lysed in 500 µL TRIzol reagent (Fisher Scientific) and stored at −80°C before purification using a standard phenol-chloroform extraction protocol with the RNAqueous Micro Kit (Thermo Fisher Scientific). RNA from three to six animals was pooled for one microarray chip. In total, four and three chips were successfully generated from muscle FAPs of young aged mice, respectively, and transcriptomic analysis was performed by a commercial provider (ATLAS Biolabs) using the Affymetrix Mouse Exon 1.0 ST Array (Thermo Fisher Scientific). Before applying statistical analysis, the input data was normalized using the quantile method and transformed to log2 values. A t-test statistic was performed and $p<0.05$ was classified as statistically significant.

ScRNA-seq was performed using two platforms. For the first analysis, FACS-purified FAPs were assessed using a microfluidics approach. FAPs ($CD45^-CD11b^-CD31^-SCA1^+$) were isolated freshly, i.e. without cultivation, from 2- and 15-months old C57BL/6J male mice at 4 dpi. Six independent chips, three per age group, employing a commercial microfluidics system (C1 System IFC, Fluidigm) were used with a capacity of capturing up to 96 individual cells per loaded chip. For each chip, FAPs were pooled from three mice (i.e. for a total of 9 mice per age group at 4 dpi). Individual cell quality was checked as per manufacturer's recommendations manually excluding doublets or low-quality cells through microscopic evaluation of captured cells, and cDNA quality control for individual cells assessed in a Bioanalyzer DNA High Sensitivity Kit (Agilent) by electropherogram tracing. Quality-filtered cells and corresponding cDNA samples of individual cells were subsequently combined on 96-well PCR plates to reduce library preparation batch effects for the “age” variable so that each plate contained 48 single cell cDNA samples from each of the two age groups, for a total of 96 samples per plate. Five such plates were processed by Smart-Seq2 cDNA library preparation (Clontech Laboratories) using the Nextera XT Indexing kit (Illumina). Each plate was then pooled into a single library, for a total of five libraries. Pooled library cDNA quality control was performed in a Bioanalyzer DNA High Sensitivity Kit (Agilent) by electropherogram tracing. Sequencing reads (single end (SE)50-mode) were generated by HiSeq-2500 (Illumina) next generation sequencing for each pool separately (one sequencing lane per pool) for a total of five (5) lanes, demultiplexed, and mapped to the murine reference genome, GRCm38 (mm10), using STAR (version 2.6.0a) with default parameters. Uniquely mapped reads were used to quantify counts using featureCounts (version 1.6.2). The R package Seurat (V3) was used for further downstream analysis, including feature selection, data integration, expression matrix scaling, and dimensionality reduction methods. Normalized counts detected expressed genes, filtering out genes expressed in $<5$ cells. Individual cell transcriptomes were excluded from downstream analyses based on the following parameters: library size $<250,000$ reads, not in G1-phase, as determined by R-based cell cycle analysis package Cyclone as described before[5], and $>10$ % reads mapped to mitochondrial genes. The gene detection threshold was set as $\geq 5$ reads in $\geq 2$ cells[6]. A total of 287 individual FAP transcriptomes remained after filtering (128 transcriptomes derived from young and 159 transcriptomes derived from aged muscle samples). Data were visualized using UMAP dimensionality reduction. Differentially expressed genes were identified using Wilcoxon Rank Sum test.

For the second scRNA-seq experiment, FACS-purified mononucleated (i.e. non-myofiber) cells were selected by Calcein blue uptake (alive cells) and the exclusion of propidium iodide (PI)-positive cells (dead cells), comparing cells of 2.5- to 15-months old female mice at 4 dpi, which were analyzed used a droplet-based single cell processing platform (10x Genomics). Approximately 8,000 sorted cells were pooled from three mice per sample to obtain heterogeneous viable cells. Cells were GEM-encapsulated using the Chromium Controller platform (PN1000202, 10x Genomics) as per manufacturers’ guidelines. Single cell-cDNA libraries were processed according to the Chromium Next GEM Single Cell 3' Kit v3.1 (PN1000269) library construction workflow, and quality-controlled using Bioanalyzer DNA High sensitivity Kit (Agilent) electropherogram tracing. Sequencing reads (paired-end (PE)75) were generated by NextSeq-500 (Illumina) NGS sequencing (28(8)56 mode) at the European Molecular Biology Laboratory’s Genomics Core Facility (Heidelberg, Germany). Data pre-processing was performed with CellRanger (10x Genomics, version

4.0.0), including filtering, barcode counting, and UMI counting, followed by alignment to reference genome, GRCm38 (mm10), using STAR (version 2.5.1b) with default parameters. The R package Seurat (version 3) was used for feature selection, data integration, expression matrix scaling, and dimensionality reduction and clustering with default parameters, unless stated otherwise. Cells with $<200$ genes expressed, and $>15\%$ reads mapped to mitochondrial genes and genes expressed in $<5$ cells were removed, resulting in 14,264 cells post-filtering (from 16,411 unfiltered cells originally). Data were visualized using UMAP dimensionality reduction. Clustering was performed with a resolution of 3. Differentially expressed genes were identified using Wilcoxon Rank Sum test. Gene Scoring[7] was applied as previously reported to identify the different cell populations using gene expression signatures described by De Micheli *et al.* [8]. Pathway analysis and gene ontology terms analyses were performed using the Metascape web-based tool[9]. $p<0.05$ and fold change $|FC| \geq 1.5$ were used as cutoff parameters.

**Data Availability of scRNA-seq datasets**

The raw data of the single-cell transcriptomic analyses reported in this manuscript have been deposited in NCBI's Gene Expression Omnibus[10] and are accessible through GEO Series accession numbers GSE247313 (droplet-based dataset) and GSE247415 (microfluidics-based dataset).

**Computational modeling of Periostin-dependent cell-cell communication**

Intercellular communication through direct cell-cell interactions (CCI) and age-related CCI-changes in the muscle stem cell niche were identified using the web-based tool scAgeCom described in[11]. Analysis was performed through filtering for the individual gene *Postn* to introduce Periostin-dependence of CCIs. The communication signals between two cell types mediated by a ligand (Periostin)-receptor interaction is based on a published dataset of limb muscle-derived scRNA-seq[11] (see also Supplementary Table 7).

**Hydrogel formation, BMDM encapsulation and immunostaining**

Modified alginate-based hydrogels with covalent crosslinking were used for encapsulation of bone-marrow derived macrophages to evaluate their polarization towards pro- vs. anti-inflammatory phenotypes in a 3-dimensional environment. First, norbornene and tetrazine were coupled to alginate as described before[12,13]. Second, norbornene-modified alginate (N-alg) and tetrazine-modified alginate (T-alg) were dissolved in PBS to a final concentration of 2% alginate. The ratio of N-alg to T-alg (N:T ratio) determined the mechanical characteristics of the material, where a ratio of 1.5 N:T was used to generate high stiffness ("stiff") materials and a ratio of 0.5 N:T was used to generate low stiffness ("soft") materials. The hydrogels were formed by mixing the N:T ratios with or without addition of 1μg/mL of mrPOSTN (R&D Systems) and a concentrated macrophage suspension. The final concentration of cells in the encapsulation was $3x10^6$ cells per mL of hydrogel. The hydrogels were left at RT for 50 minutes and then punched from the cast gel sheet using 5 mm biopsy punches (Integra 142 Miltex) and placed in BMDM differentiation medium at 37 °C and 5 % $CO_2$. After 24 hours, hydrogels with the encapsulated cells were removed from the cell culture medium and immersed in PBS for 5 minutes. Samples were then fixed in 4 % formaldehyde for 40 minutes at RT. Afterwards, the hydrogels were briefly washed with 3 % BSA in PBS and permeabilized with 0.3 % Triton X-100 in PBS for 10 minutes. Samples were blocked with 5 % BSA, 0.1 % Triton X-100 in PBS for 10 minutes at RT. The primary antibodies were rat monoclonal anti-F4/80, rabbit polyclonal anti-CD206 and rabbit monoclonal anti-CD86 (all from Abcam) antibodies diluted at 1:100 in 3 % BSA, 0.3 % Triton X-100 in PBS and incubated for 24 hours at 4°C. After incubation with the primary antibodies, hydrogels were washed three times in 3 % BSA for 5 minutes and corresponding secondary antibodies (Key Resource Table) were diluted in 3 % BSA, 0.3 % Triton X-100 in PBS and incubated for another 24 h at 4 °C. Hydrogels were washed 3 times with 3 % BSA in PBS and nuclei were counterstained with 5 μg/μl of DAPI for 30 minutes at RT. Images of three distinct areas per gel were acquired at the center region of each gel punch using a confocal microscope (TCS SP5, Leica Microsystems). For a general quantification of cell morphology, 25x magnification was used and n=6 fields of view (3 different images from 2 independent samples) were taken, containing multiple individual cells (n>50).

**Mechanical testing**

Hydrogels and tissue muscle stiffness measurements were conducted using the Test-Bench LM1 system by BOSE, along with a 250 g load cell (Model 31 Low; Honeywell). Hydrogels, after equilibration in PBS, were subjected to uniaxial unconfined compression testing at 0.016 mm/s without preload. The elastic modulus ($E$) was calculated as the slope of 2-10% of the linear region of the generated stress vs. strain curve using a Matlab script (n=4). The required Matlab inputs of hydrogel height and diameter were determined by lowering down the BOSE system top plate until contact was established or by using calipers, respectively. Tissue muscle stiffness was assessed using the elastic modulus (E). The compression step occurred with minimal pre-load, at a speed of 0.016 mm/s, extending to a maximum strain of 15% in relation to the lateral axis of the isolated tibialis anterior muscles. The elastic modulus of each muscle sample was measured once. During peak compression, the position of the device was held steady to document stress relaxation. Using ImageJ software[1], the cross-sectional area of each sample was calculated from images obtained through a digital scanner. Maximum compression was capped at 15% strain of the tissue sample's height. The elastic modulus was calculated from the slope within a 5 % range of the linear segment of the stress-strain curve.

**Human study cohorts**

Serum samples from two independent cohorts were analyzed. For one study, cohort subjects from healthy community-dwelling older (n=59; age distribution: 65-85 years; n=14 men; n=45 women) and younger adults (n=59; age distribution: 18-35 years; n=16 men; n= 43 women) were recruited between 03/2019 to 12/2019[14]. The study was registered at German Clinical Trials Register (DRKS) as DRKS00017090 and approved by the ethics committee of the University of Potsdam. Exclusion criteria were type 1 and 2 diabetes, a stroke or heart attack in the past six months, food allergies and intolerances, pregnancy or any severe or malignant disease. The group of older adults contained a sub-group of life-long athletes (n=20, 66-78 years; n=5 men; n=15 women). Older adults were assigned to this sub-group if they had participated in regular physical activity from a young age until retirement and still reported as being active at least twice a week at the time of the recruitment. Serum samples for the second cohort are part of a larger study registered at The National Center for Biotechnology Information (NCBI; clinicaltrials.gov) as NCT02994901 and has been described before[15]. In brief, patients aged ≥60 years (n=34; age distribution: 67-85 years; n=12 men; n=22 women) were consecutively recruited on hospital admission within the Department 'Geriatrics and Medical Gerontology' at the Charité - University of Medicine Berlin (recruitment was from 11/2016 to 07/2017). In addition, a healthy, age-matched control group was recruited (n=44; age distribution: 61-83 years; n=20 men; n=24 women). Exclusion criteria where neurogenerative diseases (e.g., amyotrophic lateral sclerosis or Huntington's disease), impaired cognition and the inability to understand verbal or written German. The study was approved by the Ethics Committee of the Charité - University Medicine Berlin. All participants signed a written informed consent.

**Detection of Periostin by ELISA**

Human Periostin serum levels were detected by Enzyme-linked immune-sorbent assay (ELISA; Biomedica) [16] following the manufacturer's instructions. Briefly, human serum samples were diluted 1:50 with assay buffer and incubated for 2 hours in the pre-coated microtiter plates with a mouse monoclonal anti-human antibody. After washing, samples were incubated with 150 µl of biotinylated anti-Periostin antibody for another 2 hours, washed and the conjugate was added for 1 hour. After washing the conjugate, the substrate was added for 30 minutes. The reaction was stopped and the absorbance was measured immediately at 450 nm with a reference at 630 nm using a microplate reader (Synergy H1; BioTek).

**Quantifications and statistical analyses**

All data are presented as mean ± standard error of the mean (SEM), unless specified otherwise. The sample size for each experiment and the replicate number of experiments are reported in each figure legend. Statistical significance was defined as $p<0.05$. Statistical analyses were performed using unpaired, two-tailed Student's t test or Mann-Whitney-U-test where applicable for comparison between two groups, and an ANOVA test was used for experiments involving more than two groups using GraphPad Prism software (version 9.4.1).
